# Parallel and convergent pathways for multifeature visual processing in larval zebrafish sensorimotor decision-making

**DOI:** 10.1101/2025.08.12.669772

**Authors:** Katja Slangewal, Sophie Aimon, Maxim Q. Capelle, Florian Kämpf, Heike Naumann, Herwig Baier, Krasimir Slanchev, Armin Bahl

## Abstract

Animals continuously extract and evaluate diverse sensory information from the environment to guide behavior. Yet, how neural circuits integrate multiple, potentially conflicting, inputs during decision-making remains poorly understood. Here, we use larval zebrafish to address this question, leveraging their robust optomotor response to coherent random dot motion and phototaxis towards light. We demonstrate that animals employ an additive behavioral algorithm of three visual features: motion coherence, luminance level, and changes in luminance. Using brain-wide two-photon imaging, we identify the loci of these computations, with the anterior hindbrain emerging as a multifeature sensory integration hub. Through single-cell neurotransmitter and morphological analyses of functionally identified neurons, we characterize potential connections within and across computational nodes. These experiments reveal three parallel and converging pathways, matching our behavioral results. Our study provides a mechanistic brain-wide account of how a vertebrate brain integrates multiple features to drive sensorimotor decisions, bridging the algorithmic bases of behavior and its neural implementation.

## INTRODUCTION

Animals continuously process noisy and often conflicting sensory information from their complex, dynamic environment to guide movement decisions. The precise neural mechanisms that allow the brain to extract and computationally process such cues remain poorly understood.

Multiple sensory streams can be combined through, for example, additive or winner-takes-all algorithms. For instance, when an animal attends to different targets, directing its gaze toward the averaged location can provide an effective solution^1–3^. Such an additive strategy has been observed across various species^4–8^. In experiments where animals need to combine multiple sensory features during a motor control decision-making task, additive mechanisms reliably encode signals and guide behavioral choices^9^. Alternatively, parallel representations of potential actions can compete for dominance^10^, and, ultimately, a winner-takes-all mechanism can resolve conflicting information. This competition is also thought to be fundamental to stimulus-driven attention^11^ and is widely implemented across the animal kingdom^12–16^. How these different computational strategies, adding information and winner-takes-all, are mechanistically represented across the brain and how they control behavior remains an open question.

Sensory input reaches the decision centers in the brain through different sensory modalities, like vision or olfaction, and through distinct stimulus features detected by the same sensor within one modality. Higher-order neural activity then drives a specific motor action or an adjustment of ongoing behavior. A key question is whether stimuli that cause the same behavior converge at an early stage or remain represented in parallel throughout the processing hierarchy. Early convergence reduces energy demands and enhances computational efficiency, while parallel representation allows for state-dependent behavioral flexibility. To address this topic, larval zebrafish, with their accessible nervous system, form an effective model to dissect the behavioral algorithms and neural mechanisms that underlie multifeature sensory processing within the visual modality.

Previous studies in larval zebrafish have explored the integration of separate sensory stimuli, identifying the anterior hindbrain and adjacent regions as a key site of higher-order sensory processing and behavioral control: Temporal integration of visual motion cues^17,18^, processing of heat^19^, of futility^20^, of luminance^21,22^, of internal states related to the optokinetic reflex^23^, as well as of opposing threatening looming cues^12^. In addition, this brain area controls exploratory locomotion states^24^ and represents integrated heading direction information for behavioral navigation^25^. Using a variety of visual stimuli within a single experiment, it has also been shown that stimulus representations overlap within the anterior hindbrain region^21^. However, it remains unclear how this brain area contributes to the extraction of visual features when multiple stimulus types are presented simultaneously and how it resolves situations in which sensory cues are presented in a conflicting configuration.

In this work, we disentangle how motion and luminance are integrated simultaneously in the larval zebrafish brain, combining two ethologically relevant behaviors: the optomotor response and phototaxis. Both stimuli can drive directional swimming^26–30^, and previous work has provided a good understanding of the brain regions and computations involved^17,18,21,31–35^. Our goal is to determine how these circuits interact. Are visual motion and luminance cues processed in parallel, and where do representations converge? Or are signals combined early on according to their directional influence on behavioral choice? Does a winner-takes-all competition resolve sensory conflict, or are inputs simply added to guide motor decisions? Are conflicting stimulus situations processed differently from congruent ones?

To address these questions, we first use precisely controlled behavioral experiments and construct a multifeature decision-making model. We then use our model to generate specific circuit hypotheses about the underlying neural computations in the brain. Based on such information, through brain-wide functional microscopy, we then search for respective representations. Starting at the identified loci, we then characterize potential connectivity arrangements across the computational units, allowing us to further refine our model. Thus, our study provides a comprehensive account that links behavioral experiments, neural activity and morphology, and neurotransmitter identity, enabling us to uncover a biologically plausible algorithmic and mechanistic model of multifeature decision-making in larval zebrafish.

## RESULTS

### Addition of multifeature sensory cues captures sensorimotor decision-making

Larval zebrafish swim in discrete bouts and adjust their heading direction spontaneously and based on external sensory cues. While visual motion drift and spatial luminance cues are both known to directionally guide animals, it remains unclear how zebrafish integrate these two stimuli when presented at the same time. Do they prioritize the strongest cue through a winner-takes-all strategy, or do they sum inputs across the different sensory features? To investigate this question, we systematically paired coherent random dot motion stimuli of varying strengths with either a congruent or a conflicting spatial luminance cue.

We presented visual stimuli to the bottom of 12 cm diameter arenas where unconstrained individual larvae could freely swim. A high-speed camera tracked animals in real time (**Figure 1A; Video S1**). We detected swim bouts based on increases in the variance of body orientation (**Figure 1B**), with each bout considered a decision event. To simplify our analysis, we binarized bouts based on orientation change; left (< –2°) or right (> 2°) (**Figure 1C**, top). Swim bouts occurred in stochastic interbout intervals at around 2 Hz (**Figure 1C**, bottom). To allow for variations of strength in the motion stimulus, we used random dot kinematograms of varying coherence levels..

**Figure 1.**
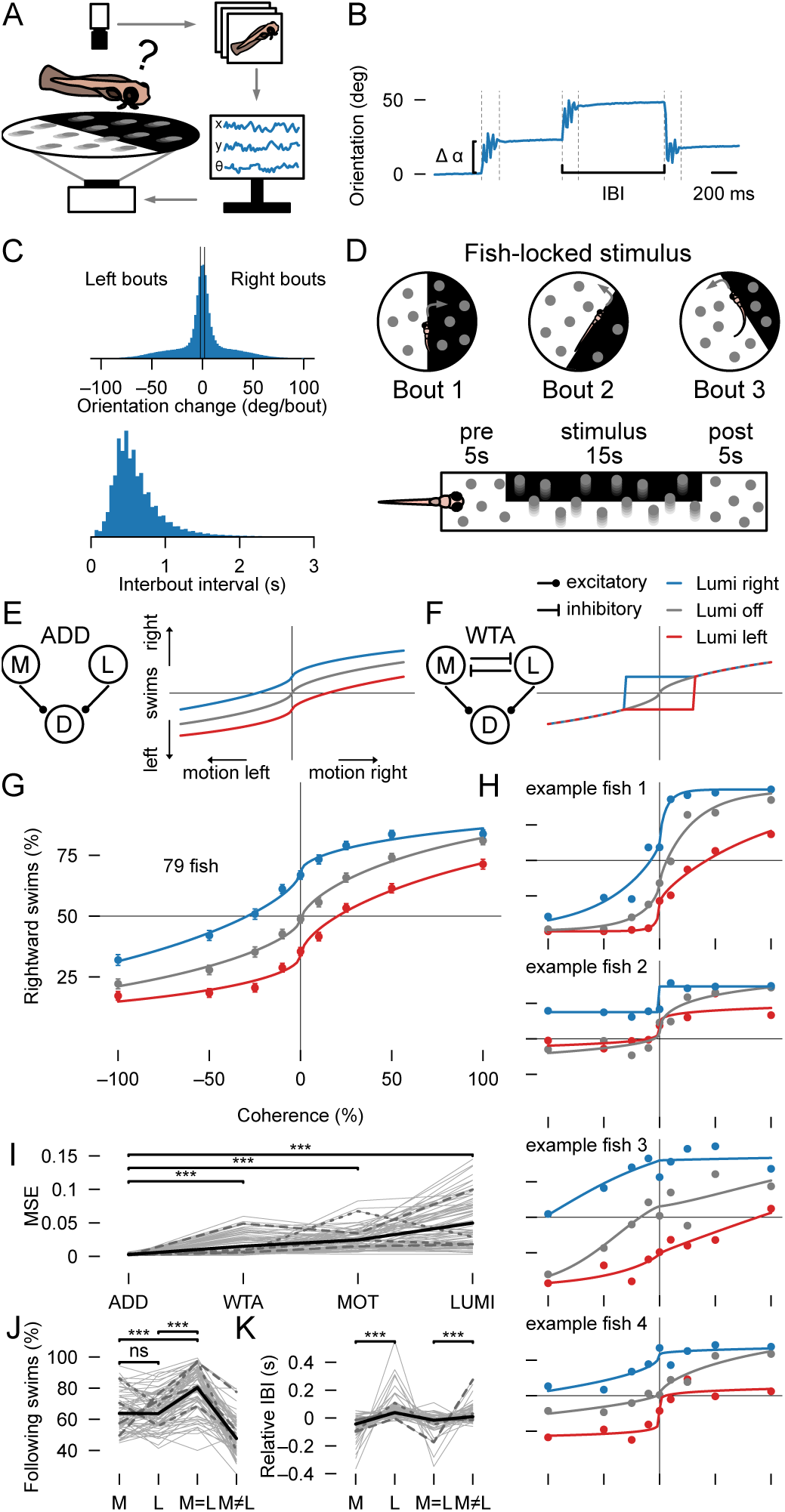
Larval zebrafish use an additive algorithm of multifeature cues to guide sensorimotor decision-making. (**A**) Freely swimming behavior setup. Individual larval zebrafish swim in a 12 cm diameter dish while being tracked with a camera at 90 Hz. Position and orientation are computed in real-time and used to update the visual stimulus projected from below at 60 Hz. (**B**) Example raw data. Δα: bout orientation change. IBI: interbout interval. (**C**) Normalized distributions of bout orientation change and IBI across all fish. We binarize bouts into left and right, ignoring straight bouts (between black vertical lines from –2 to 2 degrees) for further analysis. (**D**) Stimulus overview. We lock the stimulus to the fish position and orientation. Pre-stimulus and post-stimulus consist of 5 seconds of 0% coherence flickering dots on top of a white background. The stimulus contains moving dots at varying coherence levels, superimposed with or without lateral luminance cues. Time intervals are indicated above, stimulus order is randomly selected. (**E, F**) Expected behavioral response curves for the additive (E) and the winner-takes-all model (F). (**G**) Percentage rightward swims over coherence level. –100% means all dots move left, 100% means all dots move right. Dots and error bars indicate mean ± SEM over 79 fish. (**H**) Percentage rightward swims over coherence level for 4 individual randomly sampled example fish. (**E–H**) Axes as in (G), line colors as in the legend in (F). (**I**) Model fits of individual fish to the additive and winner-takes-all models, as well as only-motion and only-luminance configurations. (**J,K**) Percentage following swims across stimulus types (J) and relative IBI (normalized to the mean IBI across the four stimulus types calculated per fish) over stimulus type (K). M: only-motion, L: only-luminance stimuli, M=L: motion in direction of bright lateral luminance side, M≠L: motion in direction of dark lateral luminance side. N=79 fish. Double-sided t-test p-values (adjusted for multiple testing using the Bonferroni correction) are indicated with ***p<0.001, ns p>=0.05. Gray solid lines in (I–K) indicate individual fish. Dashed lines indicate the four example fish from (H) (shortest dashes: fish 1; to longest dashes: fish 4). Solid black lines indicate mean across fish. See also **Figure S1**

Certain fractions of dots moved coherently right- or leftward. All dots had short life times and got randomly repositioned in the arena, preventing animals from tracking individual dots. We superimposed coherent motion stimuli with lateral luminance cues and locked stimuli to the body orientation of animals in real-time (**Figure 1D**). This design ensures that during the stimulus, input into the visual system remains stable over time from the perspective of the fish, providing a consistent and well-controlled configuration for the study of sensorimotor decision-making. We initially focused our analysis of directional swimming to the steady-state phase within the last 10 seconds of stimulus presentation.

To uncover the computational principles underlying multifeature stimulus integration, we analyzed zebrafish behavior as a function of coherent motion strength with or without superimposed lateralized luminance cues. The behavioral decision curve in response to motion alone (= only-motion) resembles a psychometric curve (**Figure 1E,F**). What would happen to this curve if we superimpose lateral luminance cues? If zebrafish use an additive strategy, then we would expect that lateral luminance cues in which the right side is brighter than the left side would shift this curve upward. For lateral luminance cues of flipped polarity, we expect to see the curve shift downward (**Figure 1E**). If instead fish follow a winner-takes-all strategy, responses should match the only-motion curve for strong coherence levels but switch to following lateral luminance cues at low coherences (**Figure 1F**). We fitted a sigmoidal curve with additive or winner-take-all inputs^9^ (**Methods**) to make quantitative predictions for both these scenarios. Our behavioral experiments and model fitting results (**Figure 1G**) showed that, on average, zebrafish follow an additive algorithm, ruling out a winner-takes-all computation.

It may be possible that the observed effects originate from combining animals that employ distinct strategies, where some fish favor motion while others rely more on luminance. To test this idea, we fitted our multifeature model to individual fish. We fitted the additive and winter-take-all version of our model, as well as the motion-only and luminance-only versions, to account for the possibility of some individual fish relying on motion and others on luminance. While individuals exhibited variability in their response curves, the additive model described behavior better than the winner-takes-all model and single stimulus model-variations (MOT: only-motion, LUMI: only-luminance) in 76 out of the 79 tested larvae (**Figure 1H,I**). To further explore which sensory tuning parameters, such as decision-threshold and sensitivity, changes upon combining the two stimuli, we performed an additional analysis using the threshold and width parametrization of the psychometric curve^36,37^ (**Methods**). This analysis revealed that superimposing lateral luminance cues to the motion stimulus only shifts the decision threshold while leaving coding sensitivity intact (**Figure S1A-C**), further corroborating that both stimulus features are independently processed.

To further quantify behavior across individuals, we placed stimuli into four categories (**Figure 1J**, only-motion, M; only-luminance, L; congruent motion and luminance, M=L; and conflicting motion and luminance, M≠L). We picked the 25 % coherence level for the motion stimulus, since at this level, motion and lateral luminance had similar directional effects on behavior. Swims in the direction of the stimulus occurred at around 65 % in the single stimulus cases (same for M or L, as expected from our stimulus choice). Performance increased to 80 % in the congruent case (M=L) and dropped to chance levels when conflicting (M≠L), as predicted by our additive model. Interbout intervals further provided us with a proxy of reaction time and allowed us to quantify response delays for the different stimulus configurations (**Figure 1K**). We found significantly shorter interbout intervals in the only-motion compared to the only-luminance case, even though performance was the same (**Figure 1J**). The congruent and conflicting stimulus cases further enabled us to probe the algorithm of decision initiation. If motion and luminance cues would be integrated in parallel to independently trigger swim bouts, we expect interbout intervals for congruent and conflicting cases to be similar. Conversely, if motion and luminance are integrated into a common unit to then jointly control swimming, we should see shorter interbout intervals for congruent and longer interbout intervals for conflicting cases. We found significantly shorter interbout intervals for congruent than for conflicting stimuli (**Figure 1K**), supporting our latter hypothesis.

Thus, our findings demonstrate that larval zebrafish use an additive strategy to integrate coherent motion and lateral luminance cues to guide multifeature sensorimotor decision-making. We found no evidence for a winner-takes-all algorithm in this context. Furthermore, we conclude that motion and luminance cues jointly control swim initiation, leading to delayed responses in conflicting conditions. As these conclusions are based on an analysis of steady-state behavior, we next sought to examine responses on a finer temporal scale.

### A three-pathway model captures behavioral dynamics to multifeature stimulus configurations

The difference in interbout interval for motion- and luminance-driven behavior (**Figure 1K**), despite similar overall accuracy (**Figure 1J**), suggests distinct processing across the two visual stimulus features. To further explore these underlying mechanisms and expand our model across the temporal domain, we next investigated how coherent motion and lateral luminance cues are processed over time. To this end, we applied a moving average window on binarized swimming direction (**Figure 1C**, top) across trials, which allowed us to compute how the percentage of leftward swims evolves along stimulus presentation.

When fish received only-motion stimuli, they mostly swam in the direction of the motion (**Figure 2A**). The percentage of turns in this direction slowly increased upon stimulus onset and steadily decayed back to chance levels once the dots stopped moving. These findings were robust for gray dots moving on black or white backgrounds and align with previous findings of temporal integration of coherent motion cues^17,18^. Only-luminance stimuli elicited a more complex temporal response. While fish maintained a slight long-term preference for the brighter hemisphere, consistent with previous results^29,38,39^, we also observed transient peaks in response at both stimulus onset and offset (**Figure 2A**, ^40^). Notably, these peaks pointed in the direction that did not change its luminance level: At stimulus onset, fish transiently turned to the white side when the previous whole-field background was white, but to the black side when it had been black, respectively. The same was true at stimulus offset. Here, when presented with a whole-field white or black background, animals turned in the direction at which they had just seen the lateral white or black cue, respectively. When we presented motion and lateral luminance cues superimposed, we found that the behavior largely reflected the linear sum of responses to either cue alone, for both the congruent and the conflicting cases. Based on these results, we hypothesize that larval zebrafish employ at least three parallel visual pathways that process visual cues during sensorimotor decision-making: (1) temporally integrated motion (named motion pathway), (2) lateral luminance levels (named luminance level pathway), as well as (3) lateral luminance change (named luminance change pathway). Visual motion cues and bright lateral luminance levels are attractive, while sudden changes in lateral luminance are repulsive. The output of these pathways then converges without explicit mutual interactions to drive directional swimming behavior.

**Figure 2.**
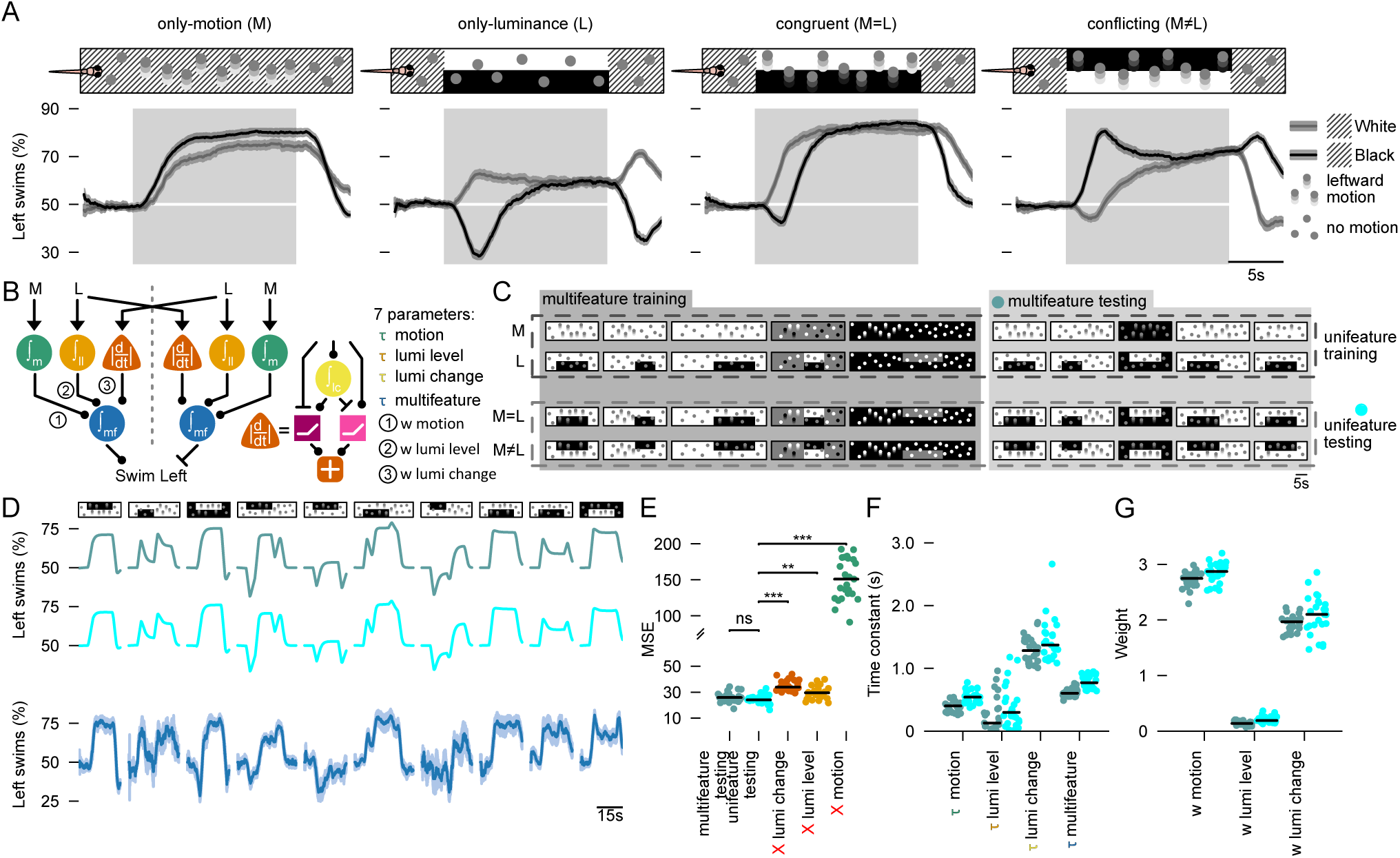
An additive three-pathway model integrating motion, luminance, and luminance change captures multifeature visual processing. (**A**) Percentage of swims to the left, over time for only-motion (M), only-luminance (L), congruent (M=L), and conflicting (M≠L) combinations. Solid gray and black lines represent trials with white and black backgrounds, respectively. Solid lines and shaded areas indicate mean ± SEM across N=64 fish. Diagonally shaded regions in the stimulus illustrations indicate whole-field white or black background periods in the absence of lateral luminance cues. (**B**) Parallel additive model with a list of the seven free parameters. Green circle node: motion integrators; golden node: lateral luminance integrator; orange triangle node: lateral luminance change detectors; blue circle node: convergence multifeature integrators. The lateral luminance change detector consists of additional units. Yellow circle: another luminance integrator. Purple square node: lateral luminance decrease detectors; pink square node: lateral luminance increase detectors. (**C**) Stimulus overview used for model testing. 40 different stimuli (**Methods** and a more detailed overview in **Figure S2A**) covering the four combination configurations of superimposed motion and lateral luminance cues. The two different model fitting approaches, multifeature and unifeature training and testing, are indicated by gray boxes and dashed rectangles, respectively. The blue and cyan dot at multifeature testing and unifeature testing indicate the labels for (D-G). (**D**) Example model predictions to 10 randomly selected stimuli not used in either multi- or unifeature training. Top row: Multifeature model prediction. Middle row: Unifeature model prediction. Bottom row: Experimental data, mean ± SEM across N=31, 13, 11, 16, 24, 16, 13, 31, 24, 11 larvae, for each stimulus, respectively. (**E**) Mean-squared-errors (MSE) for the full model (multifeature testing and unifeature testing) and for models lacking either the lateral luminance change pathway, the lateral luminance level pathway, or the motion pathway. *** p<0.001, ** p<0.01, ns p>=0.05 (two-sided t-test; p-values are adjusted for multiple testing using the Bonferroni correction). (**F**) Parameter distributions of the extracted time constants for each integrator node. (**G**) Parameter distributions of the estimated weights. The weights for the luminance level pathway were small but non-zero, matching the observed modest steady state dynamics towards lateral luminance cues in (A). Dots in (E–G) indicate individual fitting iterations of bootstrapped data (⅔ for training, ⅓ for testing, **Methods**), black line indicates median across N=25 fit iterations. Colors in (F–G) match the multi- and unifeature testing labels in (E). See also **Figure S2.**

To formalize this hypothesis, we developed an algorithmic model (**Figure 2B**) incorporating a weighted sum of these three processing pathways. The model includes seven free parameters: a weight and a time constant per pathway, and one additional time constant for a multifeature integration hub where pathways converge.

We fitted our model to experimental data and validated results through a leave-out-and-testing approach. We designed a set of 40 stimuli in which motion and luminance varied across several spatial and temporal configurations (**Figure 2C and S2A, Methods**). Stimuli classes are only-motion (M), only-luminance (L), congruent (M=L), or conflicting (M≠L). We fitted our model to the behavioral response dynamics to a subset of 20 stimuli across all four stimulus classes (multifeature training, **Figure 2C**). We then used the remaining 20 stimuli (multifeature testing) to validate the quality of the fit by computing the mean-squared error between model prediction and experimental data. Our analysis revealed that the experimental data closely matched the model prediction (**Figure 2D,E**).

We next probed whether nonlinear interactions across the three different pathways in our model may influence behavior. If no nonlinear interaction exists, then a model trained separately on only-motion and only-luminance trials should be able to faithfully predict responses to combined stimuli. In contrast, if major crosstalk occurs, then this unifeature training should result in significantly worse predictions for the combined stimuli. We trained our model on 20 stimuli selected from the only-motion (M) and only-luminance (L) classes (unifeature-training, **Figure 2C**). Notably, this unifeature-trained model performed just as well as the multifeature-trained model (**Figure 2D,E and S2B**), further corroborating the independent processing of visual information across the three parallel pathways.

To check for the relative importance of each pathway, we performed *in silico* silencing simulations. To this end, we systematically switched off individual pathways in our model (**Figure 2E**). Removing either the lateral luminance change or the lateral luminance level pathway had small, but significant, negative effects. Removing the motion integration pathway had a strong negative effect on model performance. We thus conclude that all three parts of our model are required to explain the observed behavioral dynamics.

With our fitting approach, we obtained reliable estimates of the seven parameters (**Figure 2F,G**): The time constants for the three parallel pathways and the multifeature integration hub had a similar order of magnitude, with the luminance change pathway time constant being slightly higher (**Figure 2F**). In agreement with our model unit silencing tests (**Figure 2E**) motion carried the strongest weight (**Figure 2G**). The weight for the lateral luminance level pathway played a positive, but relatively minor role. The weight for the lateral luminance change pathway was close to the weight for the motion pathway. The high weight of this pathway but the observed modest effect when silenced can be explained because the lateral luminance change pathway is only active during the short time window at stimulus transitions.

In summary, our analyses of the behavioral dynamics across the entire period of stimulus presentation support an algorithm involving the parallel integration of three visual streams: temporally integrated directional motion cues, lateral luminance levels, and lateral luminance change. While motion and bright lateral luminance cues are attractive, larvae are repelled from the side at which luminance changes occur. A simple generative model that adds and convergently integrates these processes could successfully predict behavioral response curves for a large variety of superimposed motion and lateral luminance stimuli. Having established this computational framework, we next investigated brain-wide neural activity to identify circuits that correspond to the nodes of our model.

### Model-predicted activity is represented across the brain

Since our model accurately predicted behavioral dynamics to a wide range of stimulus configurations, we next sought to determine whether and where its computational components are represented in the larval zebrafish brain. To achieve this, we performed two-photon calcium imaging in *elavl3:H2B-GCaMP8s* larvae (**Figure 3A; Video S2**). This transgenic line expresses GCaMP8s in the nucleus in almost all neurons of the brain. We presented combinations of coherent motion with and without superimposed lateral luminance cues (**Figure 3B**). By scanning different brain regions across individuals, we achieved near-complete coverage of the brain (**Figure 3C,D**).

Our model predicts the activity of seven functional nodes (**Figure 3E**): The first node temporally integrates global visual motion drift. The second one is an integrator of luminance in the luminance level pathway. Four additional nodes are found in the luminance change pathway: another integrator of luminance, followed by decrease and increase detectors and a luminance change detector that combines the two. Finally, a multifeature integrator convergently combines the three pathways. Although we can detect the general activity shape of luminance integrators, we could not distinguish between the integrators in the luminance level and luminance change pathway. Probably, H2B-GCaMP is too slow to catch subtle differences^41^, such as the 0.4 vs. 1.5 seconds time constant predicted by our model (**Figure 2F**). Hence, we treated the two luminance integrators as one unit, putting the total of functional nodes to six.

**Figure 3.**
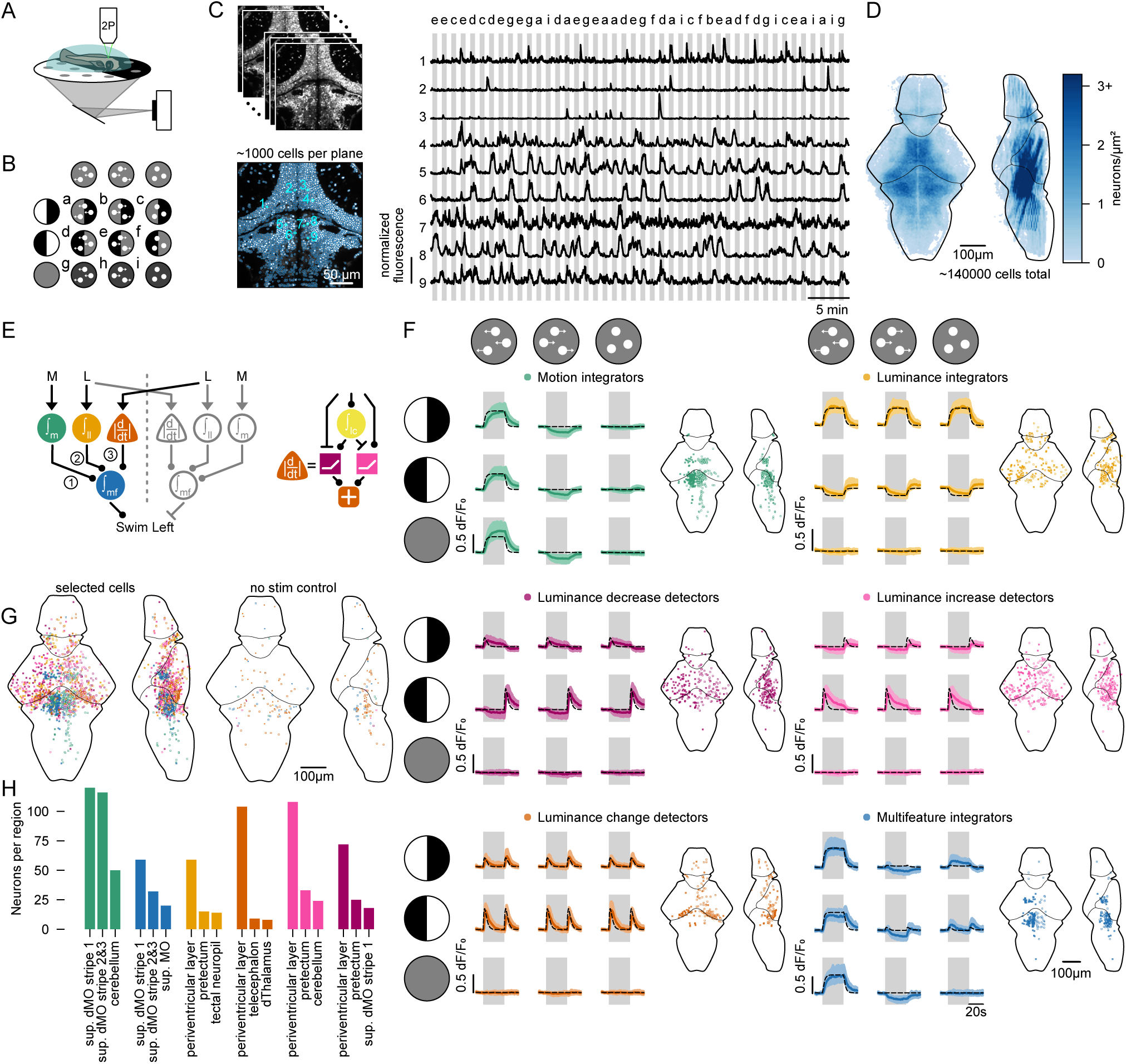
Neural representations predicted by the parallel additive model are present in the brain. (**A**) Two-photon calcium imaging of *elavl3:H2B-GCaMP8s* larval zebrafish embedded in agarose with visual stimuli presented from below. (**B**) Stimuli consisted of all combinations of motion (left, right, off) and luminance (left, right, off), resulting in 9 total combinations (a–i). Motion-off stimuli show flickering dots of 0 % coherence. Note that stimuli are illustrated in grayscale but are displayed in red to the fish during imaging (**Methods**). (**C**) Example imaging planes (top) with automated cell segmentation (bottom). Scale bar is 50 µm. Raw fluorescent example traces of 9 cells (location indicated in cyan). Y-axis scale bar indicates 1 normalized fluorescence. Letters above the traces indicate stimuli as in (B). (**D**) Overview of all imaged and segmented neurons as heatmap; the shade of blue is proportional to the number of neurons. (**E**) Model overview, as in **Figure 2B**, highlighting only the left hemisphere through solid colored circles. Colors match the legend in (F). (**F**) Neural activity of neurons found by logical statements matching each model node highlighted in (E). Solid colored lines and colored shaded areas indicate median and quartile range. Black dotted lines are the model predictions. Gray shaded regions represent periods of stimulus presentation. Before and after, we always present 0 % coherence = stimulus i in (B). Brain maps indicate the location of the found neurons. Solid circles are neurons matching the left model hemisphere, open circles are neurons matching the right hemisphere. Functional traces are combined data from the left hemisphere and flipped data from the right hemisphere. (**G**) Overview of the brain maps containing all neurons found in (F). In the control (right), we applied our logical statements to randomly selected trials of stimulus i (0 % coherence and luminance off) (see **Figure S3C** for identified control functional traces). (**H**) Number of neurons found in the five most prevalent brain regions (downloaded from the mapzebrain^42^ platform) per type from all imaged fish. N=15 fish. See also **Figures S3** and **S4**.

We detected neurons that match the model nodes using logical statements based on the temporal activation patterns predicted by our model (**Methods**). This analysis revealed a map of neural representations matching the model nodes. Specifically, we found luminance cues primarily represented in the optic tectum (**Figure 3F–H**), with additionally a high density of luminance integrators in the habenula (**Figure S3E**). Motion and multifeature integration occurred predominantly in the anterior hindbrain (**Figure 3F–H**). These representations are consistent with the projection patterns of visual inputs from the eye to the contralateral optic tectum^43^, from the eye through the eminentia thalami to the left dorsal habenula^34^ and from the pretectum to the ipsilateral anterior hindbrain^31,35^. Using a complementary classical regressor-based correlation analysis (**Methods**), we qualitatively confirmed these spatial arrangements (**Figure S3A,B**). Yet, here, we found quantitatively fewer neurons belonging to the three luminance processing units within the luminance change pathway and a wider spread of the luminance integrators (belonging to the luminance level and change pathways). These results indicate that our logical statement-based analysis can better discriminate between the different functional types than the classical linear regressor-based correlation approach.

To control for potential false-positive assignments, we applied our classifier on periods in which neither motion nor lateral luminance cues were present (**Figure 3B**, flickering dots on a dark background = stimulus i). As expected, this analysis labeled only a few neurons, without any spatial arrangement (**Figures 3G and S3C**).

Our behavioral data support an additive integration strategy over a winner-takes-all algorithm (**Figure 1E–G**). This led us to ask whether winner-takes-all dynamics may be completely absent in the brain. In the case of winner-takes-all, motion integrators should get inhibited by lateral luminance cues in a conflicting direction. Similarly, we should find luminance integrators that get inhibited when conflicting motion is shown (**Figure S3D**). Using our logical statement-based classification approach, we found such cells to be present across the brain. However, compared to neurons matching our additive model, there were fewer cells with such dynamics with less prominent spatial arrangement (compare **Figure 3F** and **Figure S3D**). These winner-takes-all cells may thus play a less important role in the sensorimotor computations under the conditions we tested in our assays. They may be recruited in different contexts, such as more complex tasks, time-constrained decisions, or at later developmental stages.

Our functional imaging results provide neural evidence that our proposed behavior-constrained model (**Figure 2B**) could indeed be mechanistically implemented on the level of the nervous system. The optic tectum emerged as a hub for detecting lateral luminance changes, while the anterior hindbrain appeared as a key site for motion integration and multifeature sensory processing. These findings suggest a structured, spatially organized circuit in which motion and luminance information are processed in parallel before being additively integrated to guide behavior.

### Orthogonal processing of motion and luminance in low-dimensional space

We next evaluated whether an unsupervised dimensionality reduction approach would yield similar conclusions to our model-based predictions. To this end, we performed a neural manifold analysis using principal component analysis (PCA) on our imaging dataset at a single-trial level (**Figure S4A**). We obtained a relatively high number of principal components: 30 components are necessary to explain 53% of the variance (**Figure S4B**). The amount of necessary components is that high, likely because our dataset aggregates neural recordings across multiple fish, with each fish imaged in different locations across the brain and being presented with a different random trial order. Nevertheless, given the constrained stimulus space of only nine combinations of motion and luminance, we hypothesized that key stimulus features would readily emerge in the leading principal components. Indeed, when projecting brain-wide activity into three principal dimensions, we identified a prominent separation between two approximately orthogonal axes corresponding to motion and lateral luminance processing (**Figure S4C**). We then computed the Euclidean distance in PCA space of opposing stimulus directions. This analysis revealed that, across the whole brain, left- and rightward stimulus conditions diverged more rapidly in the presence of superimposed luminance than in only-motion trials (**Figure S4D**). The divergence when testing congruent stimuli was greater than when testing conflicting or unifeature ones, suggesting better separability when two cues are aligned. This increased separability might play a role in the faster decision times we observed in our behavioral experiments for congruent stimuli (**Figure 1K**).

To dissect the contributions of different major brain regions to this low dimensional encoding, we separately analyzed the manifold structure in the forebrain, midbrain, and hindbrain. The luminance- and motion-related axes were present across all three major brain regions, although more strongly visible in the mid- and hindbrain (**Figure S4C**). Notably, the midbrain exhibited rapid divergence when luminance is present, suggesting that luminance changes are mostly represented in this region. The hindbrain, by comparison, showed clear separability of both motion and lateral luminance cues, with the temporal dynamics of these distance metrics largely resembling each other (**Figure S4D**).

Together, these findings revealed coarse brain-wide spatiotemporal encoding patterns for motion and lateral luminance stimuli. The unsupervised results lend convergent support to our interpretation of parallel processing and followed by convergent additive integration of motion and lateral luminance features.

### Excitation and inhibition are balanced across motion and lateral luminance processing neurons

Through our behavior experiments, algorithmic modelling, and imaging experiments, we proposed a parallel additive model of sensorimotor decision-making whose computational units are represented in the larval zebrafish brain. These experiments do, however, not allow to fully constrain the structure of model connections. For example, for lateral luminance changes on the right, an excitatory projection from the left tectum to the left anterior hindbrain would drive the fish away from these changes (**Figure 3E–H**). However, the same would be true for an inhibitory connection from the left tectum to the right anterior hindbrain. To further characterize our proposed connectivity arrangement and mode of action, we first investigated whether we find patterns in the neurotransmitter types of each functional class.

We performed hybridization chain reaction fluorescent in situ hybridization (HCR-FISH) (**Figure 4A–C**) to determine whether functionally identified neurons were (glutamatergic; vglut) or (GABAergic; gad). Knowledge about these neurotransmitter identities then allowed us to assign whether a cell would likely have an excitatory or inhibitory role during sensory processing. We aligned stacks obtained for functional imaging and post-HCR-FISH structural imaging at cellular resolution. Across all cell types, we found mostly mixed identities (**Figure 4D**). We observed a tendency for the population of luminance integrators and luminance increase detectors to be slightly more excitatory. Luminance decrease detectors seemed to be slightly more inhibitory. Notably, ratios between excitatory and inhibitory cells were largely balanced around 0.5, a common circuit motif for temporal integration^44^. We grouped cells based on neurotransmitter type, revealing no obvious differences in their functional activity (**Figure S5A**), further corroborating the idea of balanced neurotransmitter ratios. The absence of functional differences suggests these excitatory and inhibitory neurons are part of the same computation. This would indicate that the distribution of excitation and inhibition largely reflect the computations within each node, rather than signalling across nodes. To further refine our behavior-based model, we still require morphological characterization of the projection patterns across the brain.

**Figure 4.**
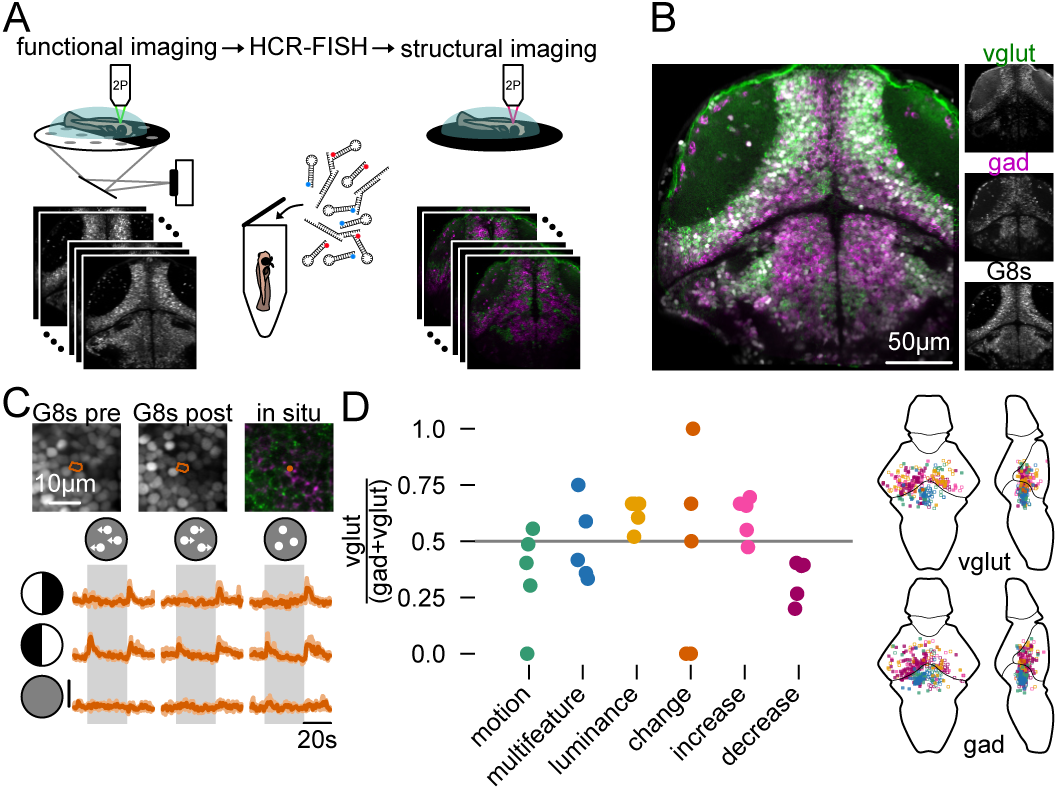
HCR-FISH reveals a balance of excitation and inhibition across computational units. (**A**) HCR-FISH pipeline with example pre-treatment H2B-GCaMP8s stack, and post-treatment gad and vglut stacks. (**B**) Overview of gad (using combined probes for gad1a, gad1b, and gad2) and vglut (using combined probes for vglut2a and vglut2b) labeling on top of H2B-GCaMP8s background in an example fish. (**C**) Example luminance change detection neuron from one example fish with its functional traces. Images on top show the H2B-GCaMP8s channel before and after HCR-FISH treatment, as well as the gad/vglut overlay. Colors as in (B). Fluorescence scale bar indicates 0.5 dF/F_0_. Solid line and shaded areas indicate median and quartile range across N=8–9 trials. (**D**) Left: Ratio of excitatory neurons (vglut/(gad + vglut)) across the six functionally identified cell types in N=5 fish. Luminance integrators had a tendency to be slightly more excitatory (t-test compared to 0.5 baseline; p = 0.15) and luminance decrease detectors a tendency to be slightly more inhibitory (t-test compared to 0.5 baseline; p = 0.17). Right: Location of the excitatory (top) and inhibitory (bottom) neurons. See also **Figure S5**.

### Neuron morphology of computational units refines model structure

Our behavioral experiments allowed us to propose a relatively simple general model structure using computational nodes. Through our imaging experiments, we found neural elements that may implement the underlying computations. Our neurotransmitter analyses revealed a largely balanced circuit arrangement. To further constrain our model based on projection patterns and to further test whether there is a brain implementation of our model, we next sought to investigate neuronal morphology. To this end, we performed two types of analyses, broadly within the identified areas as well as more precisely of the functionally labeled cell types.

First, we limited the scope of possible anatomical connections by using the morphological data available in the mapzebrain atlas^42^. We created binarized kernel density estimation (KDE) maps of the functionally identified neurons to identify relevant regions. Since these regions largely overlap between the motion and multifeature integrators, as well as for the four luminance processing types, we merged the KDE maps into one tectal map and one anterior hindbrain map (**Figure 5A**). Next, we selected neurons whose somata and projections resided in the identified regions (**Methods**). We then assigned neurons to four potential pathways through which information can flow from the tectum to the anterior hindbrain (**Figure 5B**). First, luminance information from one eye could reach the anterior hindbrain through tectal neurons projecting ipsilaterally into the neuropil ventral of the anterior hindbrain. Second, the anterior hindbrain sends ipsilateral projections anteriorly which could connect with tectal neurons who send their axons ventrally. Third, contralateral projections of the anterior hindbrain could pick up lateral luminance-related signals from ventrally projecting tectal neurons. Fourth, luminance information from one eye could reach the anterior hindbrain through tectal neurons projecting contralaterally into the neuropil ventral of the anterior hindbrain. Based on the neurons from the mapzebrain resource, each of these possibilities is feasible. However, due to the proximity of different functional cell types, it is impossible to determine if different cell types map to separate projection types, requiring a more targeted approach.

To obtain functional and structural information from the same neuron in the same animal, we performed functionally guided pa-GFP photoactivations^45,46^ (**Figure 5C, Methods**). We first selected neurons based on their functional activity. With a targeted laser pulse, we then photo-converted cytoplasmic pa-GFP, which then distributed across the whole cell, allowing us to trace the morphology of our selected neuron. Mapping reconstructed cells into a common reference frame, allowed us to compare neuronal morphologies across animals and speculate about potential connectivity based on proximity. Using this approach, we generated a library of 63 cells for which we know both their activity profile and their complete morphological structures (**Figure 5D,E; Video S3**).

We found that the multifeature neurons project either into the interpeduncular nucleus (IPN), anteriorly into the hypothalamus, or contralaterally to the other side of the anterior hindbrain, with one exception remaining local (**Figures 5E,F** and **S5A**). Most motion integrators in the anterior hindbrain project locally and remain ipsilateral, largely intermingled with the neurites of the multifeature neurons (**Figure 5E,F**). The identified luminance increase detectors all remain local in the tectum and mostly project to the stratum fibrosum et griseum superficiale (SFGS). The luminance decrease detectors also remained mostly local within the tectum, with a majority of cells projecting to the stratum album centrale (SAC) and stratum griseum centrale (SGC) (**Figure S6A**). Among the luminance change detectors we find several projections towards deeper tectal neuropil layers, the pretectum, dorsal thalamus and intermediate hypothalamus (**Figure S6A**). These ventral projections reach proximity to the anterior side of the neurites of multifeature neurons (**Figure 5G**). For the lateral luminance integrators, we largely find the same projection patterns. However, notably, projections from this cell type arrive at the posterior side of the neurites of multifeature neurons (**Figure 5E,G**).

Our combined functional and structural analyses allow us to refine our behaviorally constrained model (compare **Figures 3E** and **5H**). Our imaging experiments revealed that lateral luminance cues are processed mostly in the optic tectum on the contralateral side (**Figure 3F**). We thus propose a respective connectivity arrangement in which lateral luminance signals from the right eye get processed in the left hemisphere and vice versa. Based on our HCR-FISH results, we suggest a mixed excitatory and inhibitory balance within integrators (**Figure 4D**). This balance is likely required for the implementation of the temporal integration properties within each computational unit. Our pa-GFP photoactivation experiments show that contralateral projections of motion integrators and multifeature neurons exist. Based on our previous structure-to-function analyses of motion processing neurons in the anterior hindbrain^45^, we model these connections as likely inhibitory. We further find that the computation of lateral luminance change detection happens locally in the optic tectum. The three functional types reach the multifeature integrators through three spatially separate pathways (**Figure 5F,G**) following specific connectivity patterns: First, an excitatory connection of motion integrators onto multifeature neurons without clear compartmentalization. Second, an inhibitory ipsilateral connection from lateral luminance level integrator cells onto multifeature neurons. This arrangement is in agreement with the first proposed configuration in **Figure 5B**. Third, an excitatory ipsilateral connection from the lateral luminance change detectors onto anterior compartments of multifeature neurons. This pattern matches the second possible arrangement in **Figure 5B**. We did not find any evidence for a contralateral tectum-to-anterior hindbrain connection, making the patterns three and four in **Figure 5B** unlikely configurations. The differential targeting of the three parallel pathways onto spatially distinct compartments of multifeature neurons hints towards potential mechanisms of dendritic computations within these cells.

In summary, we have used a variety of complementary methods, involving behavioral experiments, functional imaging, HCR-FISH, and single-cell pa-GFP photoactivation to test and constrain a model of multifeature sensorimotor decision-making. The proposed model agrees with an additive strategy and has its computational nodes located in the tectum and anterior hindbrain.

**Figure 5.**
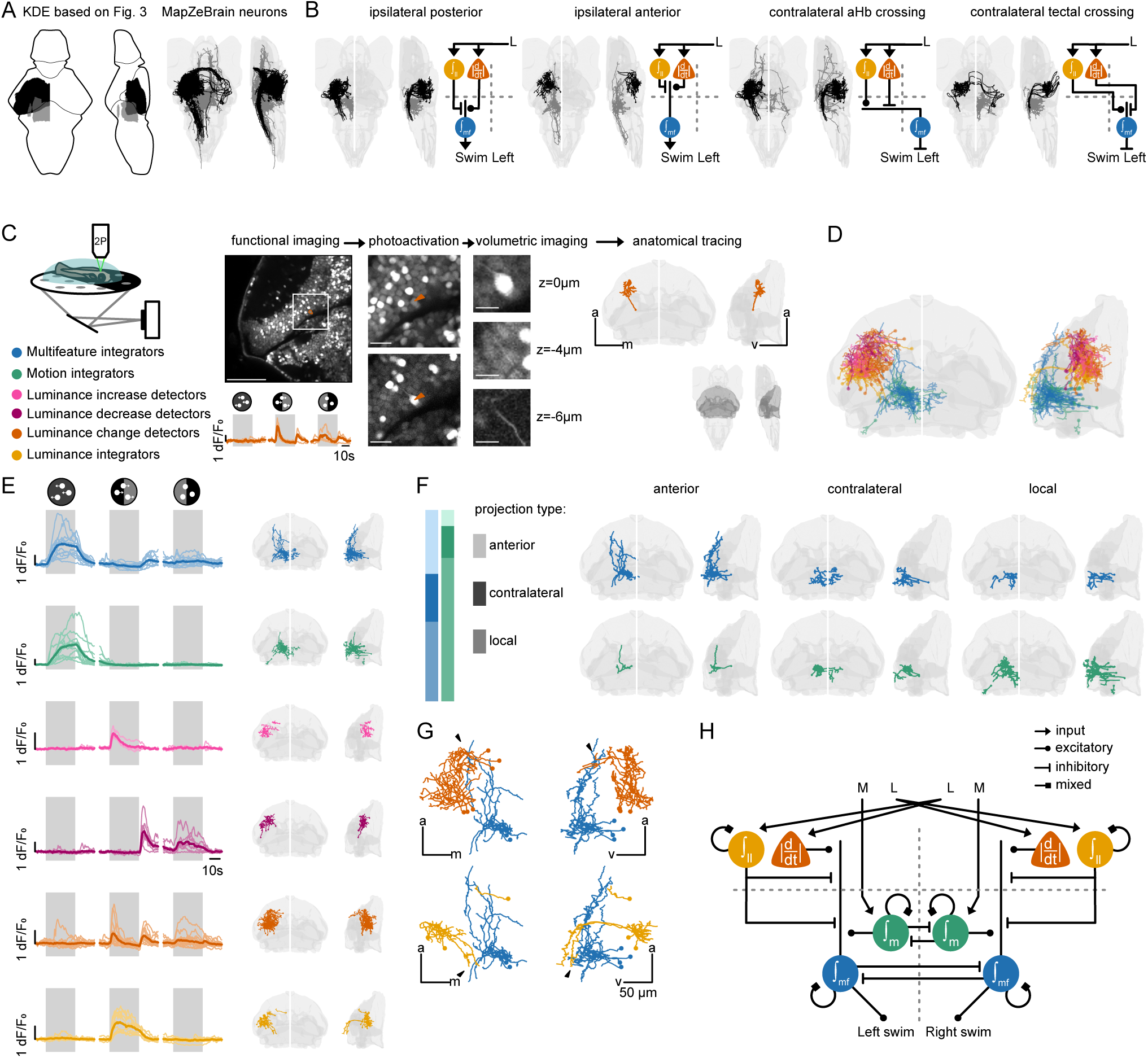
Projection patterns of functionally identified cells refine our model. (**A**) Left: binarized kernel density estimation (KDE) maps based on data in **Figure 3**, split in a tectal map (black) and anterior hindbrain map (gray). Right: all neurons found in the mapzebrain atlas within each KDE map. (**B**) Selected neurons within the identified regions, split into four potential connection patterns between tectum and anterior hindbrain. Excitatory or inhibitory type is based on the behavioral output, given the assumption that the multifeature neurons activate ipsilateral swims. (**C**) Functionally targeted pa-GFP photoactivation pipeline with an example neuron. First: functional imaging and manual cell type identification based on a minimal stimulus set containing three stimuli. Second: pre- and post-photoactivation, top and bottom, respectively. The higher-resolution image matches white square in the functional imaging plane. Third: volumetric imaging. Fourth: tracing of the photoactivated neuron. Scale bars 50 μm, 10 μm, 5 μm, respectively. Reference regions in light gray: optic tectum, anterior hindbrain sup. dMO stripe 1, and cerebellum; as highlighted in dark gray in the full brain overview. (**D**) All traced neurons of all functional types. (**E**) Functional traces and morphologies across cell types. Gray shaded area indicates when the stimulus was present. From top to bottom: 12 multifeature integrator neurons (blue), 12 motion integrator neurons (green), 4 luminance increase detector neurons (pink), 7 luminance decrease detector neurons (purple), 18 luminance change detector neurons (orange), and 10 luminance integrator neurons (yellow). Reference brain regions as in (C). (**F**) Proportion of locally, anterior, and contralateral projection of motion and multifeature neurons (left). Multifeature (blue, top) and motion integrators (green, bottom) split by projection type. (**G**) Close-up of potential connections between luminance change detectors and multifeature integrators (top) and between luminance integrators and multifeature integrators (bottom). (**H**) Refined circuit model. See also **Figure S6**.

## DISCUSSION

Our study demonstrates that larval zebrafish integrate multiple visual features—motion, luminance level, and changes in luminance—through an additive strategy to guide sensorimotor decision-making. We identify a distributed neural representation of these computations and propose a plausible circuit-level organization in which distinct stimulus features are processed in parallel and subsequently convergently combined to drive behavioral choices.

We built upon a well-established understanding of the neural algorithms underlying motion-driven behavior in zebrafish. The optomotor response has been extensively characterized as a spatial and temporal integration process involving the retina, optic tectum, pretectum, and anterior hindbrain^17,18,24,47^. Visual motion is encoded retinotopically in direction-selective layers of the tectum^48^ and pretectum^31,35,49^, and then further refined and temporally integrated in the anterior hindbrain to produce motor output^17,18^. Consistent with this framework, we observe temporally integrated motion responses in the anterior hindbrain, alongside motion-responsive neurons in both tectal and pretectal regions. Extending beyond previous work, we identify a distinct population of anterior hindbrain neurons that respond selectively to motion, and another population that integrates both motion and luminance cues.

To assess lateral luminance-guided behavior, we moved beyond the typical steady-state assessment of phototactic preference^29,50^. To this end, we adapted a paradigm in which lateral luminance cues are locked to the position and body orientation of the fish in real-time^30,39,40^. This configuration allowed us to quantify phototactic decision-making in dynamic stimulus conditions. As we could show previously^40^, phototactic behavior consists of two components: a sustained attraction to bright stimuli and a short-latency repulsion from abrupt lateral luminance changes. The neural circuitry underlying luminance processing is less well understood than the circuits for motion. A previous study has shown that varying stimulus color has differential effects on the optomotor response and phototaxis^51^. In addition, ablation of phototaxis-related retinal ganglion cells does not affect the optomotor response^50^. These results corroborate our findings of early sensory parallelization of functionally distinct visual signals. Other work on the downstream phototaxis circuitry implicates the preoptic area^52^, habenula^34^, and thalamus^50^. In our dataset, we did not find a contribution of the preoptic area, making a dominant contribution of such a pathway unlikely in our paradigm. Yet, we found luminance-responsive neurons in the habenula and thalamus, though with differing response profiles. Luminance integrators were largely present in the thalamus. Luminance integrators and luminance change detectors, but no luminance increase or decrease detectors, appeared in the habenula of some fish. While we decided to focus our analysis on the optic tectum, where all four luminance processing types were represented, we do not rule out alternative processing pathways involving the thalamus and habenula.

Previous works have employed functional lightsheet imaging to reveal luminance-sensitive neurons in the anterior hindbrain^21,22^. Results from such experiments may have been confounded by the intense blue light during imaging. Using two-photon microscopy, we confirm and expand on these findings. We find that neurons in the anterior hindbrain responding to luminance also often respond to motion as well. We conclude that these cells represent multifeature units onto which presynaptic luminance and motion-processing cells converge via an additive arrangement. This observation aligns with our behavioral experiments, in which we also showed that an additive algorithm best explains our results. Given that the anterior hindbrain and neighboring regions are involved in the processing of sensory stimuli across a broad range of modalities^17,19–21,23^, we suggest that this region serves as a hub for multifeature and multimodal sensorimotor transformations.

In agreement with previous ideas^21^, we interpret that the anterior hindbrain neurons act as an intermediate computational layer rather than directly driving motor output. This architecture introduces flexibility in behavioral control and timing. It may explain why interbout intervals are increased in conflict trials, where motion and luminance signals are incongruent. An additive algorithm to process multifeature information has been observed across species. In mammals, for example, multimodal neurons in the anterior frontal cortex^9^ and superior colliculus^53–55^ combine auditory and visual inputs using similar additive or context-modulated strategies. In humans, multisensory regions like the posterior superior temporal sulcus and middle temporal gyrus exhibit modality-specific substreams that interact depending on task context^56^.

Notably, although we observed primarily additive integration, some anterior hindbrain neurons exhibited winner-takes-all responses. These neurons may represent computational units that are selectively engaged under different task demands, developmental stages, or internal states, all of which we did not explore in this study. For example, it has been shown that a combination of averaging and winner-takes-all mechanisms govern behavior in slightly older zebrafish, when confronted with conflicting threatening looming stimuli^12^. For such directed escape responses, pure additive algorithms would blur the percept and would likely be maladaptive. Based on these reasonings, we suggest that circuit motifs for both additive and categorical decision-making coexist and can be employed under certain environmental contexts. Similar strategies have been documented in non-human primates^57^, reinforcing the idea that parallel processing supports behavioral flexibility.

Linking function and structure of the nervous system to the algorithms controlling behavior remains a fundamental and challenging goal in systems neuroscience. A major advance of our study is the integration of behavioral modeling with functional imaging and anatomical analyses, going beyond descriptive brain-wide activity maps. The types of datasets we have generated comprising neuronal function, anatomy, and neurotransmitter identity are essential for developing biophysically grounded circuit models. Recent efforts to build anatomical neuronal libraries lacking functional information^42^ have facilitated circuit dissection, as, for example, in studies of locomotion^58^, evidence accumulation^17^, retinal processing outputs^59^, and motor vigor learning^60^. It is even becoming possible to use such datasets with proximity-based metrics to speculate about brain-wide circuit motifs^61^. However, the interpretation of such analyses remains inconclusive, as functional properties and transcriptomic identity often correlate weakly^62–64^. Thus, a multifaceted approach incorporating structural and physiological data, preferably within individual behaving animals, is critically needed to understand how sensations are transformed into behavioral choices.

Here, we propose connectivity based on proximity mapping via function-guided photoactivatable GFP labeling. While spatial closeness alone does not confirm synaptic connections^65,66^, it provides a tractable and scalable framework to build testable models across individuals. Our two-photon microscopy intermediate-throughput approach balances resolution and practicality, enabling population-level modeling that can inform future targeted studies. For example, using electron microscopy, it will be interesting to explicitly search for the possible spatially confined convergence of different cell types onto multifeature neurons in the anterior hindbrain. Even without correlating function and structure in electron microscopy^45,67^, it may be sufficient to use anatomical information from existing datasets^45,68,69^ to support hypothesis testing. Over time, such a bidirectional process, moving from behavioral modeling to anatomy across light and electron microscopy datasets, and back, can iteratively refine our understanding of circuit motifs across species.

Zebrafish offer unique advantages for such studies due to the feasibility of whole-brain imaging in individual animals. Yet, similar strategies may soon be applied in mammals. For instance, functional ultrasound imaging^70^ enables brain-wide activity measurements in rodents, opening the door for cross-species comparative studies on sensory integration principles. Through optogenetic activation of superficial cortical areas in mice, it could also be shown how different sensory modalities across the visual and auditory system interact and relate to behavior in coherent and conflict trials^9^. Using such approaches in rodents, however, it remains challenging to explore behaviorally relevant computations on a cell-to-cell level.

Interestingly, our model predicted functional roles (excitatory vs. inhibitory) for certain nodes, yet neurotransmitter identity profiling mostly revealed a mixture of glutamatergic, and GABAergic neurons across all cell types. We interpret this as evidence for population-level balance, where excitatory and inhibitory inputs are combined within functionally defined nodes to shape output dynamics. Moreover, the implementation of specific temporal dynamics within each computational unit likely also requires a balanced neural network^44^. Our photoactivation results further support this view: each functional class contains locally projecting cells that could contribute to this excitation-inhibition balance. In future experiments, one can also perform neurotransmitter and anatomical analyses within the same animal and exact same neuron, allowing to further categorize cell types. In the context of motion processing, we have recently shown that such an approach could reveal inhibitory inter-hemispheric projections within the anterior hindbrain^45^.

Our anatomical analyses provide evidence for potential spatial organization of inputs onto multifeature anterior hindbrain neurons, suggesting dendritic compartmentalization, a well-known strategy for enhancing computational complexity. Dendritic segregation allows for localized synaptic integration, supporting context-dependent gating^71,72^. In flies, for example, direction-selective T4 and T5 neurons compute motion via dendritic subregion processing^73^. Similar principles have been proposed in vertebrate cortical pyramidal neurons, where distal and proximal dendrites process top-down and bottom-up signals, respectively^71,74^. Our findings suggest that zebrafish neurons may employ comparable compartmentalization to integrate visual features. Our work may thus open new possibilities to explore the computational principles and biophysical mechanisms of dendritic processing in a simple vertebrate brain.

The developed framework further lends to broader investigations into the integration mechanisms across different sensory modalities beyond the visual system. For example, combining motion with precisely controlled olfactory^75^ or thermal^76,77^ cues could allow testing whether additive integration remains the default mode or whether other rules apply across modalities. Similarly, incorporating experience-dependent changes^78^ and studying older zebrafish^40,79^ could reveal developmental tuning of integration strategies. Future work should also aim for model-based causal manipulations, though conventional ablations may lack the resolution needed for these distributed circuits in which functionally distinct neurons reside in proximity to each other. All-optical two-photon-based interrogation experiments^80^ may enable further dissections of the functional roles in the system. Emerging spatial transcriptomics approaches^50^ should further help identify molecular markers to extend the biophysical realism of our models.

Taken together, our results contribute to a broader understanding of how distinct visual features are extracted, processed in parallel, and additively integrated to guide behavior. This architecture mirrors principles of multisensory integration observed across vertebrates, suggesting that modular, parallel, and context-flexible computation is a conserved solution for transforming sensory input into action.

## Supporting information

Video S1 Behavior

Video S2 Imaging

Video S3 Photoactivations

## ACKNOWLEDGMENTS

We thank the members of the Bahl lab and Neurobiology department, as well as Joseph Donovan and Iain Couzin, for fruitful discussions. We thank Ulrike Bonitz, and our colleagues at the animal facility and the university workshop for their support. We thank Daniel Hummel, Fiona Klusmann, Ashrit Mangalwedhekar, Katrin Vogt, Joseph Donovan, and Margherita Zaupa for proofreading and feedback on this manuscript. This work was funded by the Emmy Noether Program (BA 5923/1-1), an ERC Starting Grant (101075541 – CollectiveDecisions), and the Deutsche Forschungsgemeinschaft (DFG, German Research Foundation) under Germany’s Excellence Strategy (EXC 2117 – 422037984). In addition, the Zukunftskolleg Konstanz supported A.B. and M.Q.C. The International Max Planck Research School for Quantitative Behaviour, Ecology and Evolution (IMPRS-QBEE) provided bridge funding for M.Q.C and Ka.S. Ka.S. was also supported by a Boehringer Ingelheim Fonds graduate fellowship. A.B. and F.K. were supported via the National Institutes of Health U19 Program (U19NS104653).

## AUTHOR CONTRIBUTIONS

Conceptualization: Ka.S., M.Q.C., A.B.; Data curation: Ka.S., M.Q.C., F.K., S.A.; Formal analysis: Ka.S.; Funding acquisition: A.B., Ka.S.; Investigation: Ka.S., F.K., S.A., H.N.; Methodology: Ka.S., M.Q.C., F.K., S.A., H.N.; Project administration: Ka.S., A.B.; Resources: A.B.; Software and data curation: Ka.S., M.Q.C., F.K., S.A., A.B.; Supervision: A.B.; Validation: Ka.S., M.Q.C., A.B.; Visualization: Ka.S.; Fish line generation (*elavl3:h2b-GCamp8s*): H.B., Kr.S.; Writing original draft: Ka.S., A.B.; Writing review & editing: Ka.S., S.A., M.Q.C., F.K., H.N., H.B, A.B.

## DECLARATION OF GENERATIVE AI AND AI-ASSISTED TECHNOLOGIES IN THE WRITING PROCESS

During the preparation of this manuscript, the authors used ChatGPT (GPT-4o) in order to refine readability and scientific tone. The authors then reviewed and edited the content as needed. They take full responsibility for the content of the published article.

## RESOURCE AVAILABILITY

### Lead contact

Requests for further information and resources should be directed to and will be fulfilled by Armin Bahl;

### Materials availability

This study did not generate new unique reagents. Fish lines generated in this study will be deposited to the ZFIN database upon publication (ID: ZDB-ALT-xxxxxx-xx). Fish lines will be made available upon request.

### Data and code availability

All data reported in this paper will be shared by the lead contact upon request. Analysis and model code have been deposited at GitHub and are publicly available at https://github.com/bahl-lab-konstanz/multifeature_integration_paper. Any additional information required to reanalyze the data reported in this paper is available from the lead contact upon request.

## METHOD DETAILS

### Animal care and transgenic zebrafish

We housed and handled adult and larval zebrafish (*Danio rerio*) according to standard procedures. We used the following lines: wild type AB strain, *Tg(elavl3:H2B-GCaMP8s)*^mpn438^, *Tg(alpha-tub:c3pa-GFP)*^a7437Tg^*; Tg(elavl3:Hsa.H2B-GCaMP6s)*^jf5Tg; 81^. We generated the new *Tg(elavl3:H2B-GCaMP8s)^mpn438^* line by standard procedures. In brief, we injected the Tol2 vector transgene construct *Tol2-elavl3:H2B-GCaMP8*s^82^, obtained from Janelia Research Campus, and transposase RNA into 1–4-cell-stage embryos. We then isolated transgenic lines by screening for high expression of bright green fluorescence in the central nervous system in the next generation.

We raised larvae in small groups (∼50 individuals) in Petri dishes (14.5 cm diameter) in E3 fish water + methylene blue under a 14:10 hr light-dark cycle at 28 °C. After one day, we cleaned the dishes and raised the larvae further in E3 fish water without methylene blue until 5 days-post-fertilization (dpf). We performed all experiments with larvae at 5 dpf. Sex cannot be determined at this age. Animal housing and care were approved by the Regierungspräsidium Freiburg, Germany.

### Behavior

We used WT-AB larvae for freely swimming behavior experiments, in a similar tracking setup as we developed previously^17^. In short, we transferred individual larvae into custom-designed circular acrylic dishes (12 cm in diameter, 5 mm in height, black rim, transparent base covered with diffusion paper) filled with ∼50 mL of E3 fish water. We projected visual stimuli onto the dish from below with AAXA Neo-pico projectors. Infrared LED strips (940nm panel, SOLAROX®) illuminated the dishes from below, so a high-speed CMOS camera (Basler acA2040-90um-NIR or Grasshopper3-NIR, FLIR Systems) with a zoom lens (#58-240-6X, 18-108mm FL, Edmund Optics) and an infrared filter (850nm) could record the larva.

A custom-written, Python 3.12-based software live-tracked the larvae at a rate of 90 Hz. The software determines larval position based on the largest center-of-mass after background subtraction. Next, second-order image moments define larval orientation. To identify events of high activity (bouts), the software calculates a 50-ms rolling variance window over body orientation. We defined bouts to start once the rolling window variance exceeded 1 deg^2^ for at least 20 ms. Bouts ended once the variance dropped below 0.5 deg^2^ for at least 50 ms. Besides the raw trajectories, the software stores the bout variables: time, x position, y position, and orientation at the start and end of each bout. From these variables, we determined the orientation change per bout, distance travelled per bout, and the interbout interval (time between the starts of 2 consecutive bouts).

### Visual stimuli

We used combinations of lateral luminance cues with random dot motion kinematograms. Stimuli were aligned to the position and body orientation of the fish in real-time.

Lateral luminance stimuli consisted of one bright and one dark semicircle (black and white, gray and white or black and gray) positioned on each side of the fish. The two semicircles were connected by a 0.6 cm linear transition, which prevents a distracting sharp edge directly under the fish (see also **Video S1**). For the behavior experiments, we projected white scenes at ∼1500 Lux, gray scenes at

∼900 Lux and black scenes at ∼10 Lux, unless indicated otherwise. For the imaging experiments, to prevent bleed-through into the green photomultiplier, we used red colors. We projected red scenes at ∼2 Lux, dark red scenes at ∼1 Lux and black scenes at ∼0.5 Lux. Before and after each lateral luminance stimulus presentation, we showed a baseline homogenous background. For the behavior experiments, **Figure 2C and S2A** indicate whether the homogenous background was black, gray, or white. For the imaging experiments, the homogenous background was always dark red.

On top of the lateral luminance or homogenous background, we displayed a random dot motion kinematogram. These stimuli consisted of ∼1000 gray or red dots, unless indicated otherwise (each dot was 2 mm in diameter), projected from below onto a circular arena (12 cm for behavior, 8 cm for imaging). Each dot had a mean lifetime of 200 ms and stochastically disappeared, then immediately reappeared at a random location in the arena. The coherence level, specified per experiment, indicates the percentage of dots that moved coherently to the left or right at a speed of 1.8 cm/s. We showed 0% coherence as a baseline stimulus (no movement).

During the behavior experiments, the timing of the pre-baseline, stimulus, and post-baseline varied as is indicated in the figure legends and illustrated in **Figure S2A**. We tested various combinations of motion and luminance to ensure robustness of our results. During each behavior experiment, in the first 5 seconds before the start of each actual stimulus, dots were moving towards the center of the dish to guide the fish away from the walls. This phase helped to reduce thigmotactic edge-effects. During the imaging experiments all stimuli consisted of 10 s pre-baseline, 30 s stimulus and 20 s post-baseline.

We rendered all stimuli online, using custom-written software based on Python 3.12 and Panda3D 1.10.15 with OpenGL Shading Language (GLSL) vertex shaders running on AMD Radeon RX 580 GPUs.

### Behavior data analysis

To preprocess our acquired freely swimming behavior data, we combined all bouts per fish per experiment. We excluded bouts and trials affected by tracking errors^40^. The following bouts were dropped: 1) bouts with an interbout interval > 10 s. This filters out tracking mistakes where the algorithm tracks small particles on the edge of the dish. 2) Bouts where the contour area of the fish exceeded 2000 pixels. This filters out tracking mistakes where accidental air bubbles or scratches got tracked instead of the fish. 3) Bouts with an average speed > 1 cm/s. This filters out tracking mistakes where the algorithm jumps from the edge or bubble back to the fish. 4) Bouts with an orientation change > 150°. This filters out tracking mistakes where the head and tail got swapped 5) Bouts that were within 0.25 cm or less from the edge of the dish. This removal avoided edge effects in our analysis. In addition, we dropped the entire trial, if more than 5% of the bouts were labelled as tracking errors in any of the aforementioned categories. In total this filtering step removed 13.7±11.1% of bouts and 14.3±10.1% of trials per experiment. We then binarized swim bouts into left or right bouts, ignoring all forward bouts between –2 and 2 degrees.

For the steady state behavior analysis (**Figure 1**), we calculated the percentage of leftward bouts (left / (left + right) * 100%) during the last 10 s of the stimulus. We use only the last 10 s of the stimulus since the percentage of leftward bouts reaches a steady state by then. We based the relative reaction time and following swims plots on the data from 25% coherence, since at this coherence level motion and luminance balance each other. To get the relative reaction time, we subtracted the average reaction time across stimuli (M, L, M=L and M≠L) per fish. For the percentage following swims (**Figure 1J**), we flipped the orientation data during rightward stimuli (considering motion during motion-luminance conflicts) and then merged the right and leftward stimuli.

For the analysis of decision-curves over time (**Figure 2**), we flipped the orientation data as explained above. Unless indicated otherwise, ‘percentage leftward’ plots are based on the combined data of actual leftward stimuli and flipped rightward stimuli. We then merged all trials per stimulus and calculated the percentage of leftward bouts in a rolling 2 s window (moving in 0.1 s steps). In the decision-curve plots, each window contains bouts from up to 2 s in the past.

### Additive and winner-takes-all steady-state models and fitting strategy

To determine how zebrafish integrate motion and luminance cues, we tested whether their behavior aligns with an additive model or a winner-takes-all variant of this model. Our modelling approach was based on a recent framework describing rodent multisensory decision-making^9^:

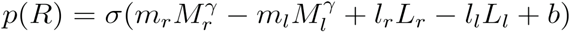

where *p(R)* is the probability of swimming right, and 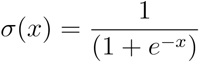 is the logistic function. Motion coherence levels are represented by *M_r_* and *M_l_* (ranging from 0 to 1), while *l_r_* and *l_l_* indicate luminance cues (either 0 or 1). In the winner-takes-all variation, the weaker stimulus is set to zero. In the single stimulus variants (MOT and LUMI) the second stimulus is set to zero. Both models include six free parameters: bias (*b*), coherence exponent (*γ*), motion sensitivity *m_r_*, *m_l_*, and luminance sensitivity (*l_r_*,*l_l_*). We fitted the percentage left over coherence curves of single fish and the group average to both the additive, winner-takes-all, and single stimulus model variants using scipy’s curve_fit function with default parameters (initial guess: 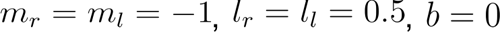, and (γ = 0.6). We used these initial guesses to create the model hypothesis plots in **Figure 1E,F**. We then use the MSE between the model fit and data to determine which model performed best (lower is better).

### Threshold-and-width-based psychometric curve fitting

To determine which sensory tuning parameter is affected by superimposing motion and lateral luminance cues, we used the threshold and width reparametrization of the psychometric curve. This version of the psychometric curve describes the tuning of the system, rather than the decision-outcomes^36,37,83,84^:

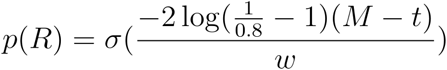

where *p(R)* is the probability of swimming right, and 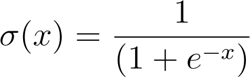 is the logistic function. Motion coherence levels are represented by *M* (ranging from –1 to 1). The two tunable parameters are threshold (=inflection point) *t*, and the width (or sensitivity) *w*. We fitted the percentage left over coherence curves of single fish and the group average to the threshold-and-width-based psychometric curve using scipy’s curve_fit function with default parameters (initial guess: *t* = 0, *w* = 0.4).

### Additive network model

Our additive network model takes four stimulus time series as input: *motion_left_*, *motion_right_*, *luminance_left_*, *luminance_right_*. All input variables are between 0 and 1 and multiplied with the clutch-specific sensitivity factors to take into account some of the variability we observe across clutches (**Figure S2A**). To simulate leaky integration of the input signals, we designed four 30-second long exponential decay kernels, with time constants 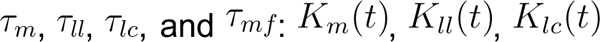, and *K_mf_(t)*:

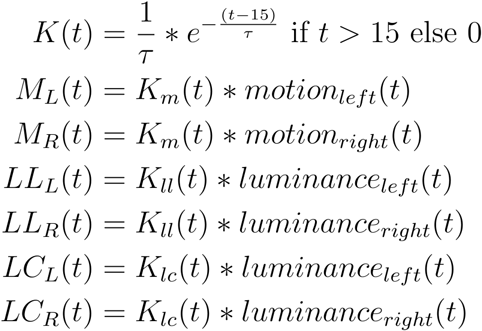

Next, we detected changes from dark to bright (luminance increase) and vice versa (luminance decrease):

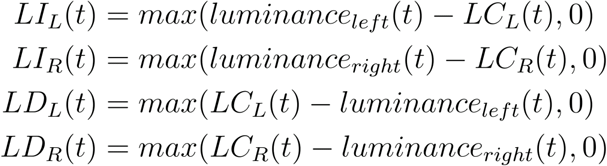

And we combined the luminance increase and decrease detectors to obtain luminance change detectors:

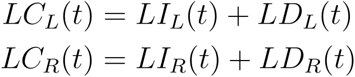

Then we took a weighted sum of the attractive luminance level and repulsive luminance change signals:

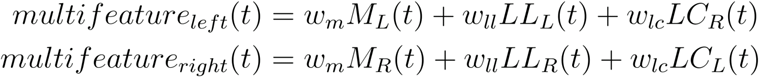

Next, we convolved the multifeature signals with *K_mf_*(t):

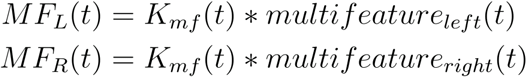

We found the ratio of left-swims:

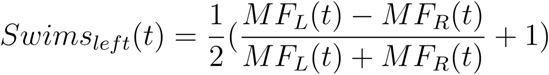

Finally, we applied a 2 second rolling average window over this ratio, to match the bout analysis.

To silence either of the three processing pathways in silico, we set either *w_m_*, *w_ll_*, or *w_lc_* to 0. Note that this also removed the respective 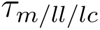 from the free parameter list.

### Model fitting

We fitted the seven free parameters 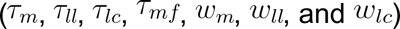 by first splitting the data 50/50 in training and validation groups. Either we put all unifeature trials (motion OR luminance) in the training group and all combinations (motion AND luminance) in the validation group, or we put half the experiments across all trial types in the training group and the other half in the validation group.

During each of the 25 training rounds, we further split the training group into training and test sets. Two thirds of the fish were part of the training set, one third of the fish were part of the test set. We then picked five times 20 random combinations of each trial kind and experiment type. For each of the five times, we set the boundaries and randomly picked the initial guess of the seven fitting parameters(*T_x_* in [0.1, 100], *w_x_* in [0.1, 25]). Using Scipy’s curve_fit() function we fitted the additive network model to the training data set and obtained a MSE score by comparing the fitted model to the test data set. We stored the best out of five parameter sets for validation, meaning that we ended up with 25 parameter estimations. Testing these 25 parameter sets on the validation group gave us 25 validation MSE scores. We used these MSE scores to show whether unifeature training of the model was sufficient to explain motion/luminance combination trials. Finally, we used the median of each fitted parameter for visualization of the model node activities in **Figure 3**.

### Two-photon imaging

We screened *Tg(elavl3:H2B-GCaMP8s)* and *Tg(alpha-tub:c3pa-GFP; elevl3:H2B-GCaMP6s)* larvae for GCaMP expression. At least 1 h prior to imaging we embedded each larva within a 6 cm diameter Petri dish in low melting agarose ∼2% (Ultra Pure Low Melting Point Agarose, 16520-100, Invitrogen) in E3 fish water.

After embedding we transferred the fish under one of our custom-built two-photon microscopes. We controlled the microscopes through custom-written Python 3.12-based software (PyZebraPhysiology). In short, our microscopes consist of a shared tunable DeepSee MaiTai Ti:Sapphire laser (SpectraPhysics) operated at 950 nm to image GCaMP and tuned to 760 nm to photoactivate c3pa-GFP. We regulated the laser power with a combination of a lambda-half plate (Thorlabs, AHWP05M-980) and a Glan-Thomson prism (Thorlabs, GL5-A) to 12–15 mW at sample for functional imaging, and 5–7 mW at sample for c3pa-GFP activation. We scanned with a set of x/y galvanometers (Cambridge technology). One microscope had a 20x NA 1.0 (XLUMPLFLN Olympus) objective, the other a 25x NA 1.1 (CFI75 Apo 25XC W 1300, Nikon) objective. We collected emitted light through GaAsP (green channel) and Alkali (red channel) photomultipliers (Hamamatsu) and amplified the signals with a current preamplifier (TIA60, Thorlabs). We presented visual stimuli through a mirror by a P300 Neo Pico Projector (Aaxa Technologies) onto diffusive paper glued to the bottom of the experimental platform (8 cm diameter).

We scanned the functional imaging planes at ∼1 Hz and 800x800 pixels at a resolution of 0.20 to 0.65μm per pixel. The vertical distance between planes ranged from 2 to 12μm. We imaged each plane for an hour and showed a random combination of nine different 1-minute stimuli, meaning that on average each stimulus was presented 6–7 times. We imaged 1–40 planes per fish. After each imaging session, we collected one overview stack (∼50 planes, 800x800 pixels at a resolution of 0.65 x 0.65 x 2 μm) to allow anatomical registration to the ZBRAIN and/or mapzebrain atlas^42,85^.

### Imaging preprocessing

We preprocessed the raw imaging data using a custom-written Python 3.10-based preprocessing script. In brief, we performed CaImAn piecewise rigid motion-correction^86^ followed by Cellpose automatic segmentation^87,88^. For the functional experiments and initial photoactivation experiments the following parameters were used: model_type=’cyto’, diameter=12, cellpose_flow_threshold=0.95. For the rest of the photoactivation experiments the following parameters were used: model_type=’cyto3’, cellpose_flow_threshold=0.4, cellpose_prob_threshold=0. Whe then used scipy.interp1d for stimulus temporal alignment^89^ (discretization=0.5 s), followed by a two-step ANTs anatomical registration to the ZBRAIN/mapzebrain atlases^42,85,90^ (step 1=overview to atlas mapping, step 2 = functional plane to overview mapping). We used the following ANTs mapping parameters: ‘interpolation_method’: ‘linear’, ‘use-histogram-matching’: 0, ‘matching_metric’: ‘MSE’, ‘rigid’: { “t”: “Rigid[0.1]”, “m”: “MI[$1,$2,1,32,Regular,0.25]”, “c”: “[1000x500x250x300,1e-8,10]”, “s”: “3x2x1x0”, “f”: “8x4x2x1”}, ‘affine’: {”t”: “Affine[0.1]”, “m”: “MI[$1,$2,1,32,Regular,0.25]”, “c”: “[200x200x200x100,1e-8,10]”, “s”: “3x2x1x0”, “f”: “8x4x2x1”}, ‘SyN’: {”t”: “SyN[0.1,6,0]”, “m”: “CC[$1,$2,1,2]”, “c”: “[200x200x200x100,1e-7,10]”, “s”: “4x3x2x1”, “f”: “12x8x4x2”}.

We normalized fluorescence 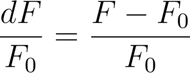 with *F_0_* being the average fluorescence during the pre-stimulus period (0 to 10 s).

### Imaging analysis linear regression

We used the model predictions of the six nodes as regressors after convolving them with a GCaMP response kernel, an exponential decay with a time constant of 2.4 s (this corresponds to a half decay time of 3.5 s^41^). We then used scipy’s linregress function with default parameters to find the correlation coefficient between each regressor and the normalized average trace of each neuron. For the integrator nodes (motion, luminance, and multifeature), we used a threshold of 0.85, for the detector nodes (increase, decrease, and change) we used a threshold of 0.65.

### Imaging analysis logical statements

To select which neurons fitted with the model predictions, we first summarized the response of each neuron into five values: A) Average pre-stimulus dF/F_0_ (0 to 10s), B) Average initial stimulus dF/F0 (12.5 to 17.5s), C) Average stimulus dF/F0 (20 to 40s), D) Average initial post-stimulus (42.5 to 47.5), and E) Average post-stimulus (50 to 60s). We then used a set of logical statements to select our neurons of interest. M=Motion stimulus, L=Lateral luminance cue.

Motion integrators:

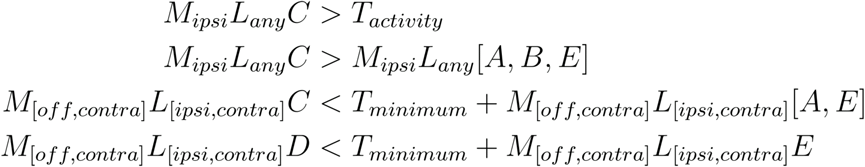

Multifeature integrators:

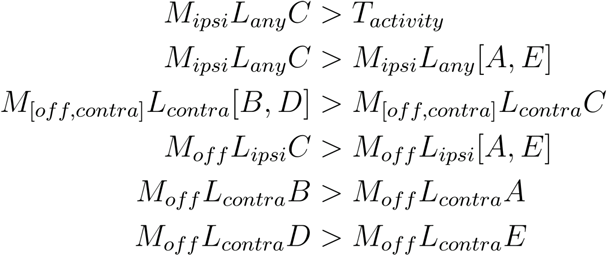

Luminance integrators:

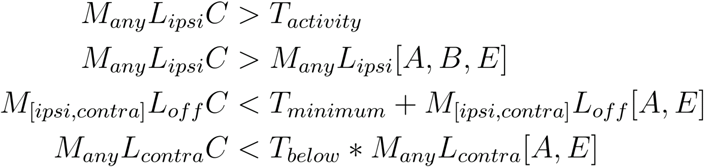

Luminance change detectors:

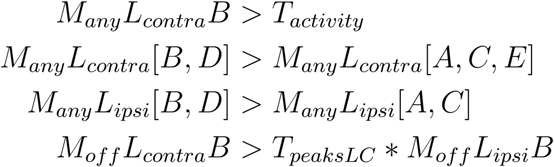

Luminance increase detectors:

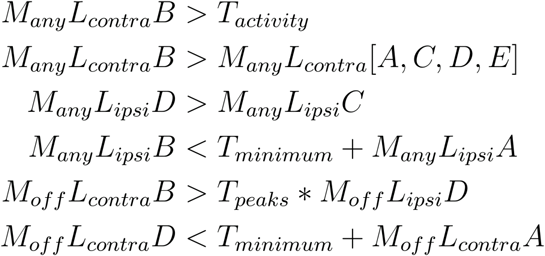

Luminance decrease detectors:

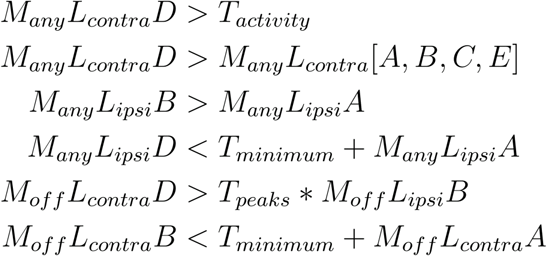

We then selected the neurons using thresholds: *T_activity_* = 0.2, *T_minimum_* = 0.1, *T_peaksLC_* = 1.25, *T_peaks_* = 1.5, *T_below_* = 0.9.

### Neural manifold analysis

We performed our neural manifold analysis across all neurons that had seen at least 3 trials per stimulus. Due to the random stimulus order there were a few planes where this was not the case, we excluded the neurons on those planes (∼30.000 neurons out of ∼140.000 were excluded).

We normalized all raw fluorescent traces and stacked the stimuli ordered as in **Figure 3B**. We performed principal component analysis (PCA) using sklearn.decomposition’s PCA function on either all neurons, or the neurons from a specific major brain region (Forebrain, Midbrain, Hindbrain). We visualized the first three principal components (PCs) labeled by stimulus type (only-motion, only-luminance, congruent, and conflicting) or by stimulus direction (right and left).

Next, we calculated the Euclidean distance in 3-dimensional PCA space between left and right trials of the same type over stimulus time, as well as the control distance between trials of the same type and same direction.

### KDE binary mask generation and mapzebrain atlas-based anatomical analysis

To create the kernel density estimation (KDE) binary masks, we took the neurons per functional class as selected by our logical statements. We took the centroid locations and used the scipy’s stats.gaussian_kde functionality with default parameters. We created a downsampled grid in ZBRAIN^85^ coordinate space (10x downsampled for computational time reasons). We evaluated the density of this downsampled grid by putting it through the computed KDE and upsampled 10x to get to the original ZBRAIN coordinates and resolution. Finally, we binarized our mask by setting every pixel within either optic tectum or superior dorsal medulla oblongata stripe 1-3 in the left hemisphere with a density > 2^-7^ to True. Next, we mapped the ZBRAIN coordinate space to mapzebrain coordinate space using a custom generated bridge transform (available upon request). Because of the high anatomical overlap, we took the union of the motion and multifeature masks, as well as the union of the four luminance related masks. We then searched the mapzebrain atlas for neurons that had their soma in either of the two masks.

For each anterior hindbrain neuron we then checked whether it crossed the hemisphere (x > 311 in ZBRAIN units), whether it projected anterior (y < 500 ZBRAIN units), posterior (y > 780 ZBRAIN units), ventral (z < 30 ZBRAIN units), dorsal (z > 90 ZBRAIN units), and/or lateral (x > 400 ZBRAIN units). These thresholds were selected with the aim to separate projection patterns without having to manually evaluate each single neuron. We defined anterior_ipsi neurons to project anterior, but not dorsal nor ventral nor cross to the other hemisphere. Anterior_contra neurons project anterior, to the other hemisphere and lateral. Local_ipsi neurons do not project anterior, posterior, dorsal or cross the hemisphere. Local_contra neurons cross the hemisphere, but do not project anterior, posterior or dorsal.

For each tectal neuron, we checked whether it projected entirely within the tectum, whether it crossed the hemisphere (x > 311 in ZBRAIN units), projected anterior (y < 400 ZBRAIN units), posterior (y > 780 ZBRAIN units), ventral (z < 50 ZBRAIN units), and/or dorsal (z > 120 ZBRAIN units). We define front_pathway neurons as projecting anterior, outside the tectum, but not posterior nor crossing the hemisphere. Lateral_pathway neurons project ventral outside the tectum, but not anterior nor posterior. Front_crossing neurons project anterior, outside the tectum, but not dorsal nor posterior. Posterior_pathway neurons project posterior outside the tectum.

In **Figure 5B** we plotted the local_ipsi anterior hindbrain neurons with the lateral_pathway tectal neurons; the anterior_ipsi anterior hindbrain neurons with the front_pathway tectal neurons; the anterior_contra anterior hindbrain neurons with the lateral_pathway tectal neurons; and the local_ipsi anterior hindbrain neurons with the lateral_cross_neurons.

### Photoactivations and morphological analyses

We performed functionally targeted photoactivations using a double-transgenic *Tg(alpha-tub:c3pa-GFP; elevl3:H2B-GCaMP6s)* line. As we have done previously^45^, we outcrossed this line to Tg(alpha-tub:c3pa-GFP), generating a high likelihood of offspring to be homozygous for *alpha-tub:c3pa-GFP* but heterozygous for *H2B-GCaMP6s.* Compared to other strategies using neuronal promoters and a single-construct design (FuGIMA)^35^, our approach helped to strongly increase c3pa-GFP expression strength, while keeping GCaMP background fluorescence at lower levels. We embedded the fish as described above at least 2 h prior to the experiment. First, we checked for expression of c3pa-GFP by photoactivating an interneuron in the right hemisphere of the tectal neuropil. These neurons project locally, so their photoactivation did not interfere with our later tracings of neurons in the left hemisphere.

We then functionally imaged 1 to 4 planes of interest for 9 minutes each, showing 3 repetitions of motion-left-luminance-off, motion-right-luminance-right, motion-off-luminance-left stimuli. These three stimuli allowed us to distinguish between all predicted cell types. We imaged at 950 nm, 8–10mW, at a rate of ∼1Hz, with a pixel resolution of 0.21 x 0.21 μm. We manually selected our cells of interest using FIJIs^91^ plot-z-axis-profile-live tool or using our custom-made Python 3.12-based software.

We photoactivated the selected neuron by performing 1 to 4 rounds of 20x 200ms-ON-100ms-OFF pulses of 760 nm 6–8mW light, focused on the center of the selected cell.

To prevent motion artifacts while imaging the photoactivated neuron, we anesthetized the fish with 300 μL of 0.015% MS-222 solution (Sigma-Aldrich, E10521-50G) directly after the photoactivation. While waiting for the MS-222 to diffuse through the agarose, we imaged a close-up stack of the photoactivated neuron (950 nm, 7–8mW, 800x800 pixels, 30–40 planes, 0.084 x 0.084 x 0.5 μm resolution). This high resolution imaging allowed us to verify whether we truly hit a single neuron and none of its neighbors. About 20 minutes after the photoactivation, we imaged a broad volume around the neuron of interest (950 nm, 8–12mW, 800x800 pixels, 60–100 planes, 0.65 x 0.65 x 2 μm resolution, 90 s average per plane). We initially traced a single neuron per fish. To increase efficiency, we later identified up to 8 additional neurons per fish, which we photoactivated and scanned after acquiring the broad volume of the single neuron.

We traced the photoactivated neurons in FIJI using the SNT tool^92^, resulting in SWC files for each traced neuron. We registered these SWC files to the ZBRAIN/mapzebrain^42,85^ atlas and extracted the brain regions (mapzebrain database downloaded on 19-06-2025 from the official website https://mapzebrain.org) through which each neuron passed.

### HCR-FISH

We performed functional experiments as described above, using the same 9 stimuli (at least 3 trials per stimuli), across at least 3 planes. The two-photon excitation wavelength was 950 nm, pixel lateral resolution of 0.3 μm, and frame rate was 1.3 Hz. We also recorded the volume using higher axial resolution (2 μm) to help registration with the post-in situ volume.

Larvae were fixed in ice-cold 4% paraformaldehyde (PFA) immediately after functional experiments, dehydrated in methanol, rehydrated, hybridized overnight at 37 °C with DNA probes, and fluorophore-conjugated hairpins were then added and amplified via hybridization chain reaction (HCR) according to standard procedures^93,94^. Probe sets and reagents were obtained from Molecular Instruments. We combined B1-*gad2*, B1-*gad1a*, B1-*gad1b*, B2-*slct17a6a* (vglut2b), B2-*slc17a6b* (vglut2a) probes, and used B1-546 and B2-405 as hairpins.

We then used the same two-photon system again to image HCR-FISH-labeled volumes. To detect the 546 nm red fluorophore, we imaged at an excitation wavelength of 1010 nm, simultaneously using two PMTs: one to detect green fluorescence from H2B-GCaMP8s and the other to detect red fluorescence from the HCR RNA-FISH probe. For detection of the 405 nm blue fluorophore, we imaged each z-plane at two excitation wavelengths. First, we used 950 nm excitation to image H2B-GCaMP8s with the first PMT. We then switched to 800 nm excitation to detect the blue fluorophore using the second PMT. In all configurations, laser power was 12–20 mW and planes were imaged for 30 to 120 s. Volumes were recorded at pixel lateral resolution of 0.3 microns, and axial resolution of 2 μm.

### HCR-FISH analysis

In situ imaging stacks were mapped to functional volume stacks using Ants^90^ with these parameters: ‘rigid’: {”use”: True, “t”: “Rigid[0.1]”,”m”: “MI[$1,$2,1,32,Regular,0.25]”, “c”: “[1000x500x250x300,1e-8,10]”, “s”: “3x2x1x0”,”f”: “8x4x2x1”}, ‘affine’: {”use”: True, “t”: “Affine[0.1]”, “m”: “MI[$1,$2,1,32,Regular,0.25]”, “c”: “[1000x500x250x100,1e-8,10]”, “s”: “3x2x1x0”, “f”: “8x4x2x1”}, ‘SyN’: {”use”: True,”t”: “SyN[0.1,6,0]”, “m”: “CC[$1,$2,1,2]”, “c”: “[200x200x200x100,1e-7,10]”, “s”: “4x3x2x1”,”f”: “12x8x4x2”}.

Because GABAergic and glutamatergic neurons can be interspersed and located directly adjacent to each other, and because alignment between the functional imaging volume and the post-fixation in situ-labeled volume can be imperfect, automatic assignment of neurotransmitter identity would be error-prone. To ensure accurate classification, we used a custom-written GUI to manually inspect the alignment and assess gad and vglut expression for each neuron individually. Only neurons that could be identified as GABAergic or glutamatergic with high certainty were included in the analysis. Neurons were excluded if alignment was uncertain, if too few surrounding features were available to assess registration, if the labeling was too noisy, or if gad and vglut signals overlapped too strongly to allow clear discrimination. This led us to keep 315 out of the 2240 model-relevant neurons identified in these experiments for which neurotransmitter assignment confidence was very high.

### Statistics

Figure 1: Two-sided t-test for related samples to compare MSE between models ADD, WTA, MOT, and LUMI, we used Bonferroni correction for multiple tests. Cohen’s D effect sizes; ADD vs. WTA: – 1.17, ADD vs. MOT: –1.96, ADD vs. LUMI: –1.80. Two-sided t-test for related samples to compare relative interbout-interval, we used Bonferroni correction for multiple tests. Cohen’s D effect sizes; M vs. L: –1.57, M=L vs M≠L: -0.84. Two-sided t-test for related samples to compare following-swims distributions, we used Bonferroni correction for multiple tests. Cohen’s D effect size: M vs. L: –0.035, M vs M=L: –1.07, L vs M=L: –1.08. Figure 2: two-sided t-test for independent samples to compare the MSE distributions, we used Bonferroni correction for multiple tests. Cohen’s D effect size: multifeature vs. unifeature full model: 0.39, full model vs silenced luminance change pathway: -2.58, full model vs. silenced luminance level pathway: –1.09, full model vs silenced motion pathway: -5.60. Figure 4: two-sided t-test for independent samples between the excitatory and inhibitory ratio and 0.5 obtained no significant results. **Figure S4**: two-sided t-test for independent samples between mean distances during stimulus time for congruent and conflicting trials. Cohen’s D effect size: 2.36 full brain, 2.74 rhombencephalon. Video legends S1 to S3

## SUPPLEMENTARY MATERIAL

**Figure S1.**
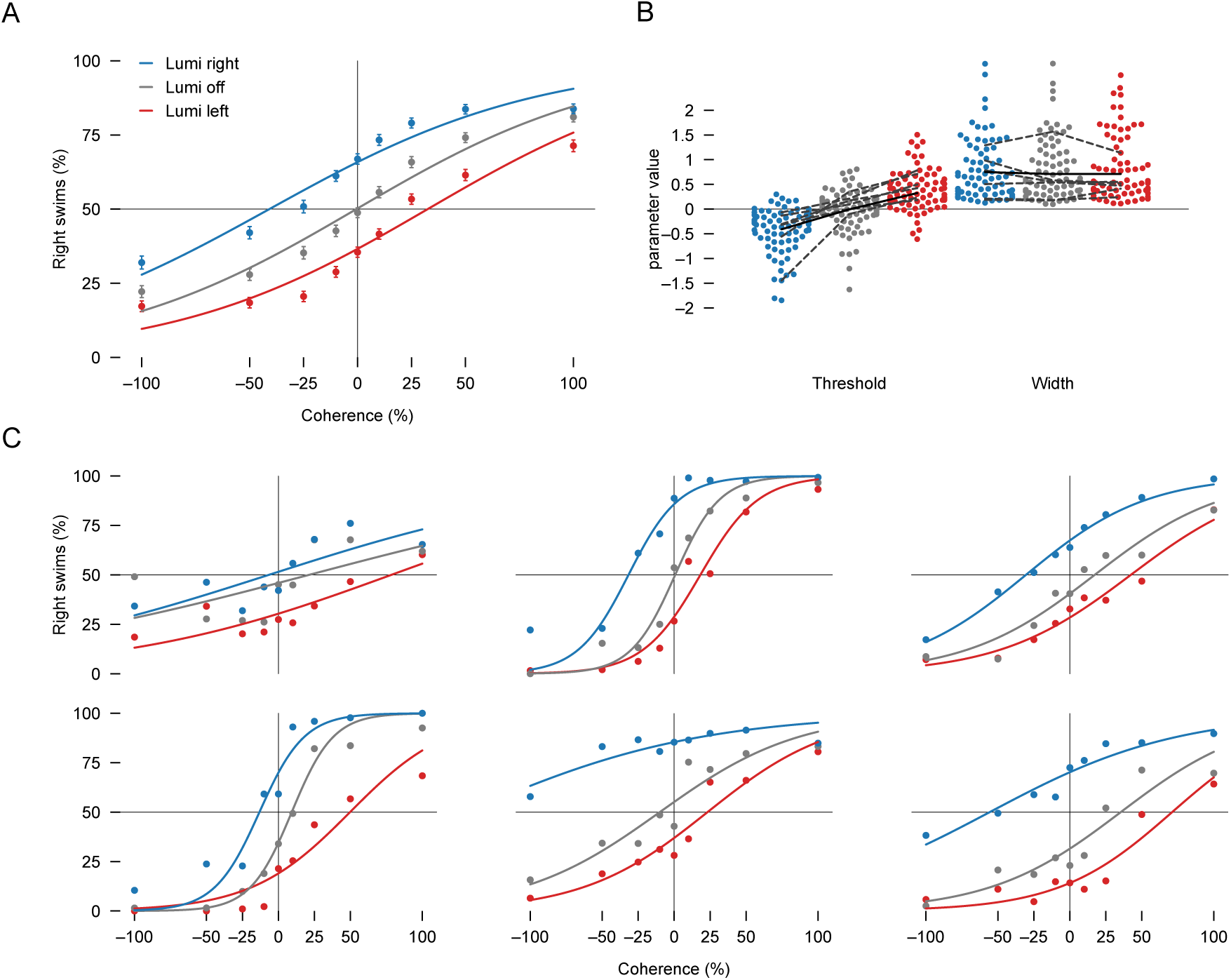
Fitting of threshold and width based psychometric curves reveals changes in threshold rather than width when combining motion and luminance stimuli. (A), Same data as in Figure 1G, but fitted with a threshold-and-width-based psychometric curve, also known as the threshold and support parametrization^36,37^. (B) Threshold and width parameter estimates for each individual fish (small filled dots). The threshold indicates how much the inflection point is shifted across the x-axis, the width indicates the coding region that contains the 10–90% response rate. Solid black lines are the fit to the mean across all fish. Dashed gray lines belong to the six example fish in (C). (C) Six randomly picked example individual fish and their psychometric curve fit. Related to Figure 1.

**Figure S2.**
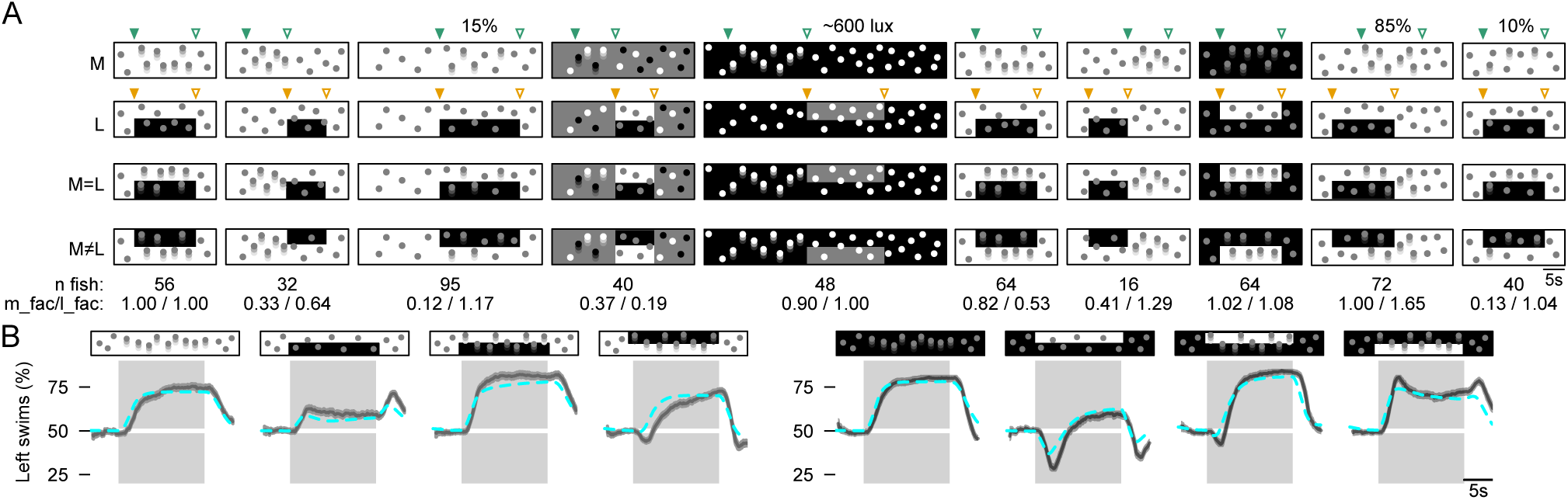
Stimulus detail overview. (A) Overview of the 10 stimuli used for the behavioral modelling experiments. Green arrowheads (top row) indicate start (solid) and end (open) of motion. Yellow arrowheads (second row) indicate start (solid) and stop (open) of the lateral luminance stimulus. Motion coherence was 50% unless indicated otherwise in the top row. White luminance was ∼1500 lux, black luminance was ∼10 lux, gray luminance was ∼900 lux unless indicated otherwise in the top row. The bottom row shows the number of fish per stimulus set, as well as the motion_factor and luminance_factor that was fitted to the data. (B) Model fit to the behavioral data shown in Figure 2A. Gray lines and shaded curves indicate mean ± SEM (same data as in Figure 2A), cyan dashed lines indicate model fit, gray shaded blocks indicate the time when the stimulus was on. Related to Figure 2.

**Figure S3.**
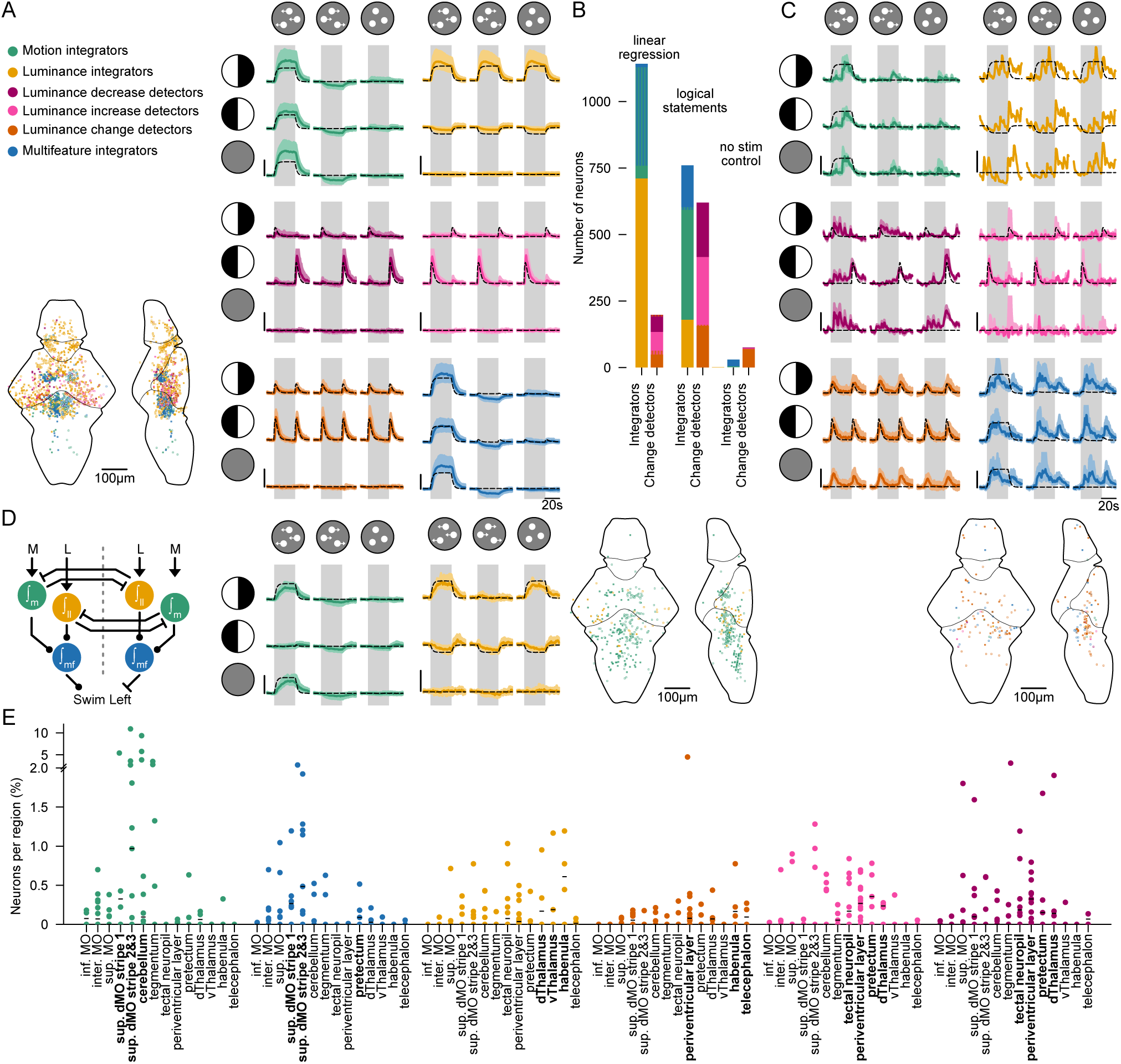
Linear regression separates functional classes less well than logical statements, control analysis of imaging data reveals no spatial structure, and winner-takes-all representations exist but are spatially distributed. (**A**) Neural activity of neurons found by linear regressor-based correlation to each model node (**Figure 3E**). (**B**) Number of neurons found by linear regression (left), logical statements (middle), and control analysis (right). Striped blocks indicate neurons that fall into two categories. (**C**) Control analysis where we applied our logical statements to randomly selected trials of stimulus i (0 % coherence and luminance off). (**D**) Neural activity of neurons found by logical statements matching motion and luminance integrators who interact through a winner-takes-all mechanism (illustrated in model cartoon). Colors and brain illustrations in (A, C, D) as in **Figure 3F,G**. (**E**) Percentage of neuronal cell type per region (as compared to the total number of neurons imaged in that region). Each dot represents a fish. Data is only considered when we imaged at least 100 neurons in the region per fish. Black lines indicate median percentage. Related to **Figure 3**.

**Figure S4.**
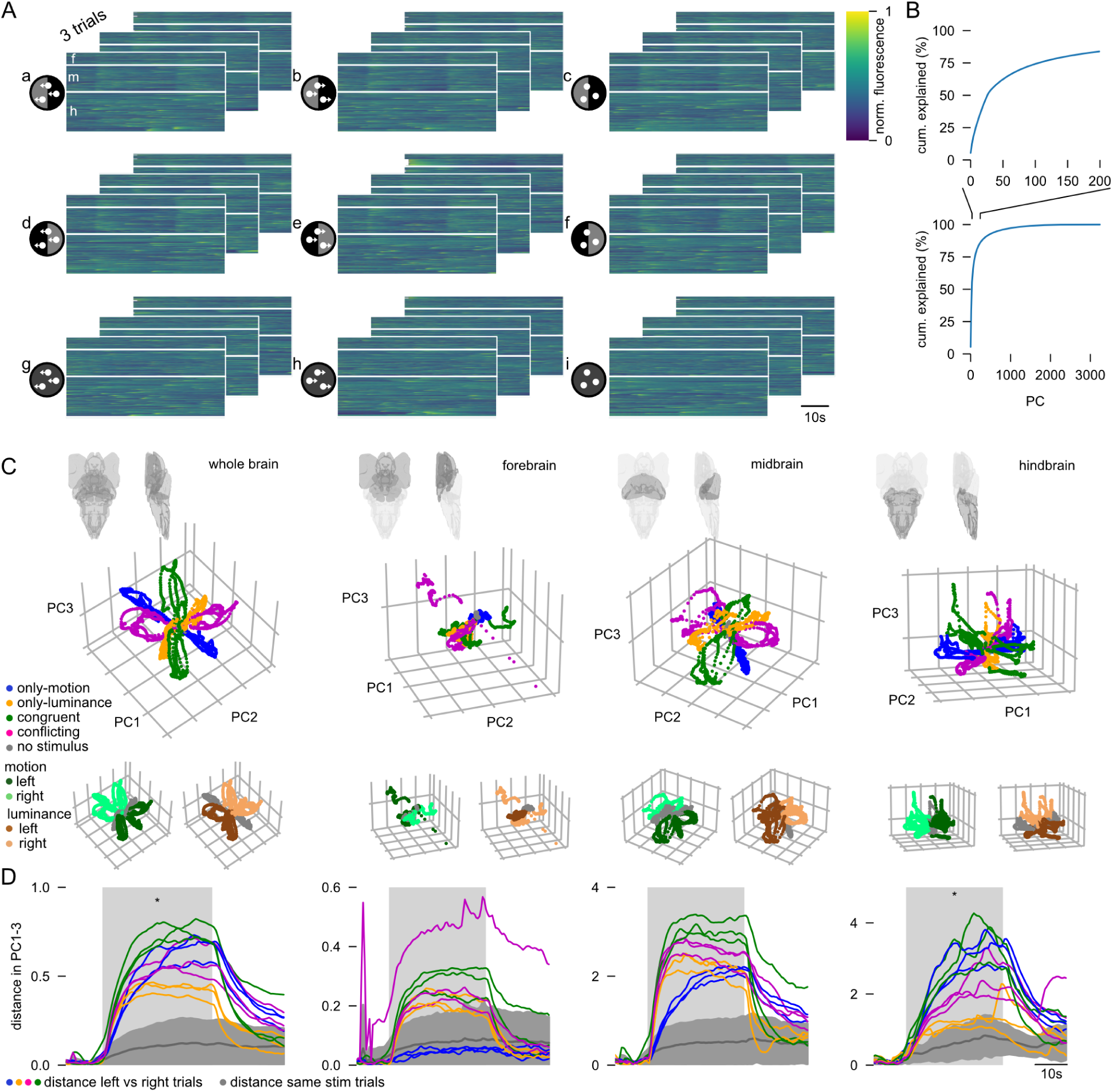
Unsupervised neural manifold analysis reveals orthogonal motion and luminance encoding with different timescales across brain regions. (A) Overview of the raw data: functional responses of 108118 neurons to 9 stimuli across 3 trials from N=15 fish (same animals as in **Figure 3**). Traces are ordered by major brain region: f-forebrain, m-midbrain, h-hindbrain. (B) Cumulative explained variance across principal components (PCs). Top: zoom-in of full range in bottom plot. (C) Neural manifold in PCs 1 to 3 for the entire dataset and subsets of the forebrain, midbrain, and hindbrain. The manifold is labeled by stimulus type (only-motion, only-luminance, congruent, conflicting), and below by stimulus direction (left, right). (D) Distance in low dimensional space between two trials of the same type but opposite direction across stimulus time. Gray line and shaded area show the mean ± STD of the distance between 2 trials of the same type in the same direction. Light-gray shaded box indicates period of stimulus presentation. Stars indicate significant difference in mean distance over stimulus time (light-gray shaded box) between congruent and conflicting trials according to a two-sided t-test: * p<0.05. Figure related to **Figure 3**.

**Figure S5.**
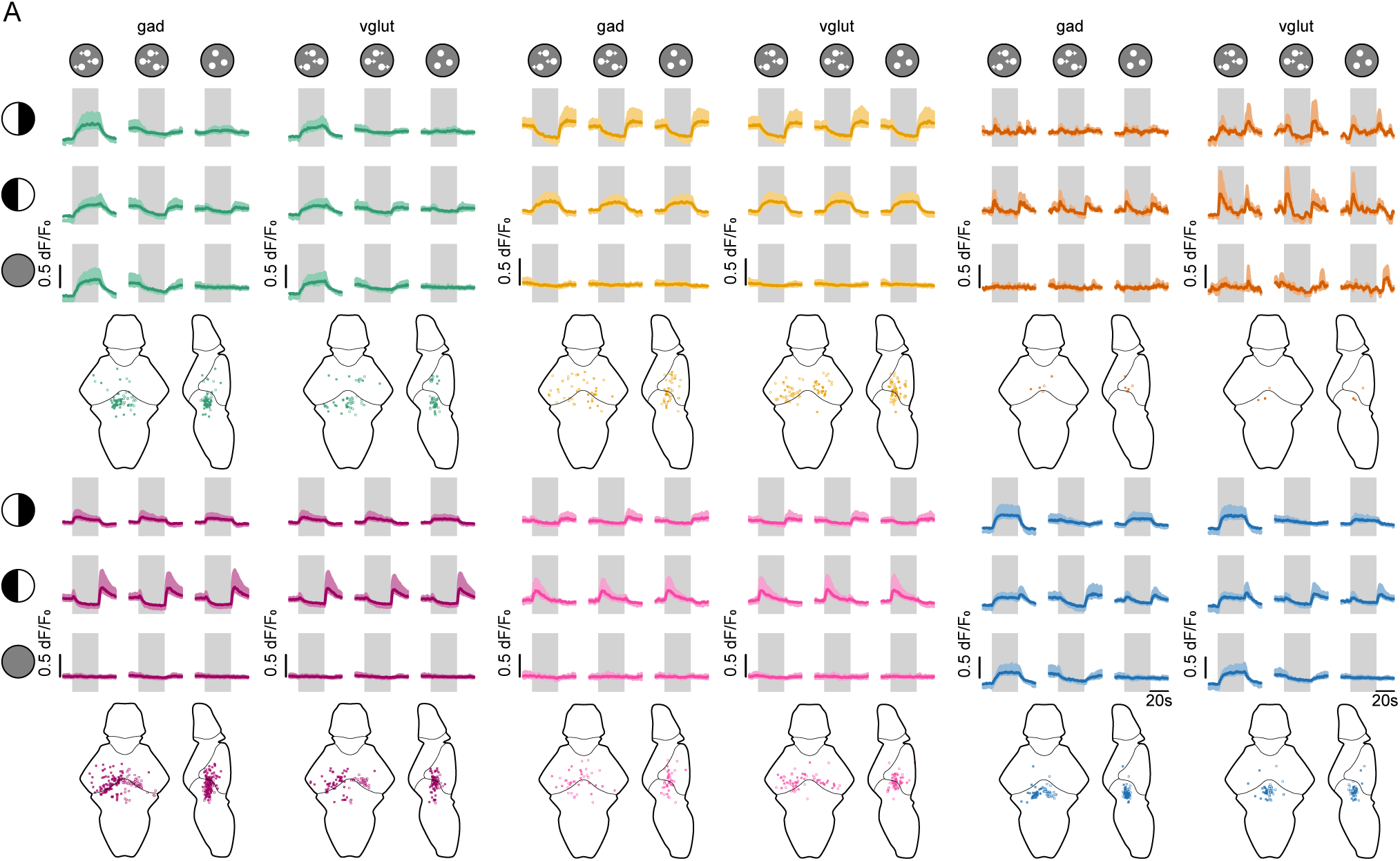
Gad- and vglut-positive neurons show no clear differences in their functional activity patterns. (**A**) Activity traces and location for each functional cell type split by neurotransmitter type. Left columns: gad; Right columns: vglut. Top row from left to right: motion integrators, luminance integrators, luminance change detectors. Bottom row from left to right: luminance decrease detectors, luminance increase detectors, multifeature integrators. Activity traces and brainmaps as in **Figure 3F**. N=5 fish with 3–9 trials per stimulus. Related to **Figure 4**.

**Figure S6.**
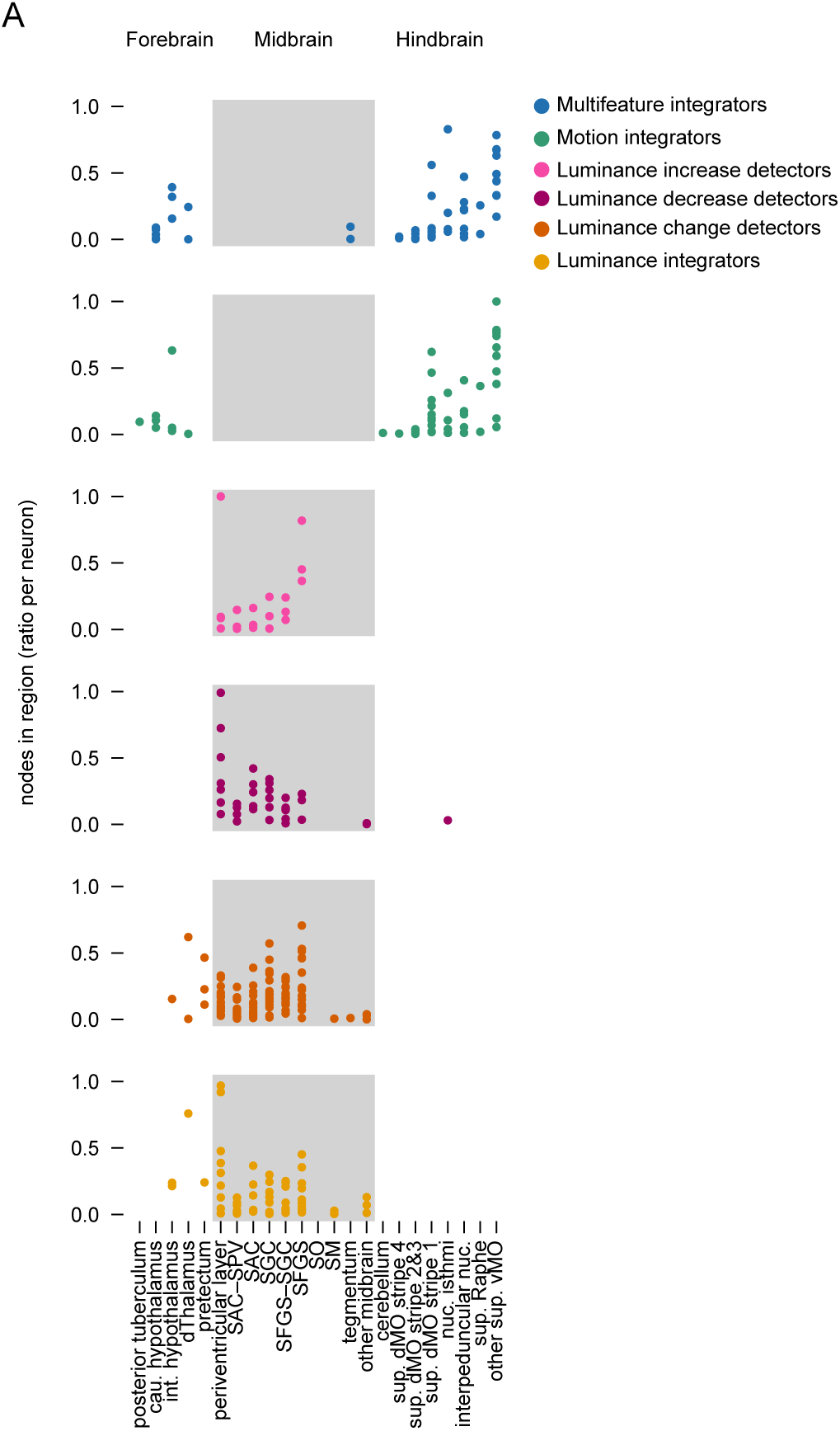
Projection patterns of photoactivated neurons. (**A**) Ratio of nodes in a brain region per neuron. Each dot represents one neuron. Data is only shown if the neuron is present in the region. Abbreviations: cau - caudal, int - intermediate, d - dorsal, SAC - stratum album centrale, SGC - stratum griseum centrale, SFCS - stratum fibrosum et griseum superficiale, SO - stratum opticum, SM - stratum marginale, sup - superior, MO - medulla oblongata, nuc- nucleus, v - ventral. Regions are ordered by forebrain, midbrain, hindbrain. Gray background area distinguishes these three major regions. Figure related to **Figure 5**.

## Video legends S1 to S3

**Video S1. Behavior examples.** One example fish, freely swimming in a 12 cm diameter circular arena. Visual stimulus consisting of coherent dot motion and/or lateral luminance cues. The video shows in order: motion stimulus, luminance stimulus, congruent stimulus, conflicting stimulus. Each stimulus consists of 10 s pre-stimulus baseline, 30 s stimulus, 10 s post-stimulus baseline. The video is displayed at 2x the original speed. The blue trace was added post-hoc to visualize the recent trajectory of the fish (750 frames; ∼8 seconds into the past).

**Video S2. Imaging examples.** Trial-averaged neural activity of three example imaging planes from three different fish (forebrain - top left; midbrain - bottom left; hindbrain - bottom right). The video shows in order: motion stimulus, luminance stimulus, congruent stimulus, conflicting stimulus as indicated by the visual stimulus in the top right. Each stimulus consists of 10 s pre-stimulus baseline, 30 s stimulus, 20 s post-stimulus baseline. The neural data videos are displayed at 5x the original speed, the stimulus illustration runs at original speed. Scale bar is 50 μm.

**Video S3. Photoactivated neurons.** Three-dimensional view of the photoactivated neurons in a rotating reference brain. Green: motion integrators, yellow: luminance integrators, pink: luminance increase detectors, purple: luminance decrease detectors, orange: luminance change detectors, blue: multifeature integrators.

